# Pneumolysin nanopores with a 20 nm diameter characterize the size and shape of individual Tau oligomers

**DOI:** 10.1101/2025.02.07.637128

**Authors:** Anasua Mukhopadhyay, Wachara Chanakul, Yu-Noel Larpin, Saurabh Awasthi, Anna D. Protopopova, René Köffel, Alessandro Ianiro, Michael Mayer

**Author notes:** Corresponding Author Email ID.

## Abstract

Protein nanopores enable the characterization of single proteins in solution without labeling by monitoring the pore’s electrical resistance when protein analytes enter their lumen. A long-standing challenge with biological nanopores, however, is that proteins in their folded conformation are often too large to enter these pores. Here, we report the self-assembly of 35 ± 5 monomers of the pneumolysin (PLY) toxin into stable transmembrane pores with diameters of 20.2 ± 3.0 nm, effective lengths of 9.5 nm, and excellent low noise characteristics in the context of nanopore-based resistive pulse recordings. The exceptionally large diameter of PLY pores enables the characterization of the size and shape of individual proteins, protein complexes, and amyloid oligomers with molecular weights ranging from 50 to 660 kDa. Using PLY nanopores, we followed Tau protein oligomerization in solution by revealing molecular weight, aggregation number, approximate shape, and oligomer concentration over time. At least four characteristics make PLY nanopores attractive for the analysis of heterogeneous amyloid oligomer samples; they: 1) provide stable open pore current with high signal-to-noise ratio; 2) enable label-free analyses of single oligomers without clogging; 3) detect a range of amyloid oligomer sizes, shapes, and aggregation numbers; and 4) reveal the concentration of each oligomer subpopulation.

## Introduction

Measurements of ion current through a single nanoscale pore embedded in an insulating membrane enable rapid, label-free detection and characterization of individual macromolecules.^1–7^ Nanopores fall into two main categories: solid-state nanopores, fabricated from inorganic materials,^7–9^ and biological nanopores, formed by self-assembling protein domains or subunits in lipid bilayers.^10–13^ In a typical nanopore experiment, analytes are added to a compartment containing high-salt recording buffer to provide a stable baseline current through the nanopore (open pore current). Entry of a non-conducting analyte particle into the pore typically causes a measurable reduction of the ionic current, called a resistive pulse. The amplitude of current blockade and the modulation of the intra-event current during this pulse provide quantitative information about the size and the shape of the analyte molecules.^1–3,8,14,15^

In the past three decades, several biological nanopores, including α-hemolysin (α-HL), *Mycobacterium smegmatis* porin A (MspA), and *Curli secretion* protein G (CsgG), have provided exciting advances in the context of DNA sequencing because their constriction diameter of 1–2 nm is suitable for translocating and interrogating the different nucleobases in linearized DNA or RNA strands.^13,15–17^ Biological nanopores, such as α-HL and Fragaceatoxin C (FracC), have been used to detect unfolded proteins, peptides, and polymers.^10,12,18^ Quantification and characterization of proteins in their native folded state were achieved, using either biological nanopores and relatively small analyte proteins or solid-state nanopores for large proteins and protein aggregates.^19–21^ These studies provided information about protein conformation and function,^22–29^ formation of protein complexes,^11,30–33^ misfolding of proteins,^34,35^ aggregation of proteins,^36–40^ biomarker detection,^26,28,41^ and introduced novel approaches for diagnostics and therapeutic monitoring.^42–44^

Biological nanopores have several advantages compared to solid-state nanopores: they can provide lower electrical noise,^45^ they are not prone to pore clogging by non-specific absorption of the analyte to the pore walls,^11,12^ and they often provide translocation speeds of analytes through the pore lumen that are slower than the translocation speeds through most solid-state nanopores.^12^ Yet, analysis of mid-sized to large folded proteins with biological nanopores requires pores with inner diameters larger than the analytes of interest. Some of the largest biological nanopores available to date for the investigation of single folded proteins are two-component pleurotolysin pore, PlyAB, with a *cis* pore opening of 10.5 nm, and a trans pore opening of 7.2 nm, and an internal constriction of 5.5 nm,^20,21^ the *Yersinia enterocolitica* pore-forming toxin, YaxAB, with a *cis* opening of 15 nm, and a *trans* constriction of 3.5 nm,^46^ and complement component 9, Poly(C9), with a cylindrical pore diameter of 10 nm.^19^ These nanopores provide information on analyte proteins with a maximum size between 125-230 kDa,^19,46^ this range is too small for the detection of large protein complexes and aggregates, such as clinically relevant amyloid oligomers of Tau proteins that are implicated in neurodegenerative diseases.^47–50^

To introduce a biological nanopore suitable for the analysis of protein assemblies with sizes approaching the megadalton range, we introduce here the use of Pneumolysin (PLY), a cholesterol-dependent bacterial cytolysin^51^ that belongs to the family of pore-forming toxins (PFTs).^52^ Cryogenic electron microscopy (CryoEM) revealed that 42 monomers of PLY self-assemble into a cylindrical pore with an outer diameter of 40 nm, an inner diameter of 25 nm, and a pore length of 11 nm (**Figure 1a**).^51^ Both the large size and cylindrical shape of this pore make it an attractive candidate for use in nanopore experiments with natively folded protein analytes.

**Figure 1.**
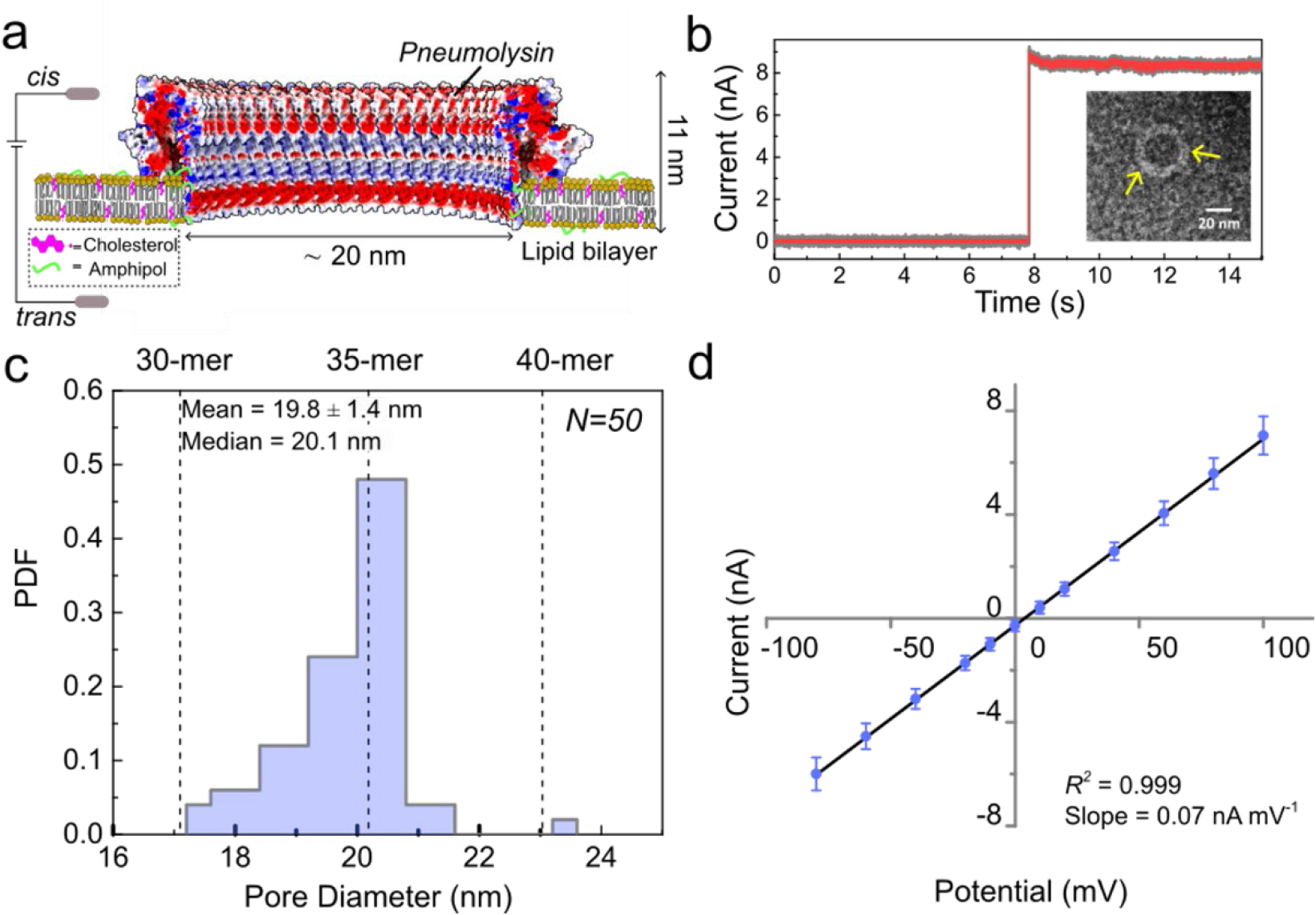
Characterization of self-assembled PLY nanopores in planar lipid bilayer membranes. (**a**) Cartoon with a surface representation of a section of a PLY nanopore in a cholesterol-containing planar lipid bilayer (PDB 5AOE). The lumen of the PLY nanopore is colored by electrostatic potential, with red indicating negative charge and blue indicating positive, as computed by the adaptive Poisson–Boltzmann solver (APBS).^71^ (**b**) Current Recording showing a single-step insertion of a single PLY nanopore into a planar lipid bilayer under +100 mV applied potential to the *trans* compartment (the *cis* compartment is connected to electrical ground). The data shown in grey was sampled at 200 kHz with no filter applied, and the data shown in red was filtered with a digital Gaussian low-pass filter with a cutoff frequency of 10 kHz. The recording buffer contained 500 mM NaCl, 50 mM Tris-HCl pH 7.5, and 0.02 mg mL^-1^ amphipol. The inset shows a TEM micrograph of the ring-shaped PLY assembly formed after incubation of PLY monomers in an aqueous solution containing amphipol, in the absence of lipids. (**c**) Distribution of PLY pore diameters calculated assuming a cylindrical pore shape^72^ from single-channel conductance measurements of PLY pores in cholesterol-containing planar lipid bilayers under an applied potential of +100 mV, using an effective pore length of 9.5 nm (see **Supplementary Note 2** for a discussion about the effects of deviations from a perfectly circular cross-section of PLY pores).^51^ The top axis in (c) represents the estimated number of PLY monomers that corresponds to the determined values of pore diameters (see **Supplementary Note 1**). (**d**) Current-voltage (I–V) curve through a PLY pore under the same conditions as in (b). Error bars represent the standard deviations calculated from at least three repeats.

To demonstrate the applicability of PLY nanopores to the analysis of large protein assemblies, we explored the assembly of microtubule-associated protein Tau into oligomers.^53–55^ Full-length human Tau protein (Tau441(P301L), 45.9 kDa) is an intrinsically disordered protein with multiple cellular functions, including transport of neuromodulators.^56,57^ Aggregates of Tau, often in the form of hyperphosphorylated Tau protein, are a characteristic pathological feature in a wide range of neurodegenerative diseases, including Alzheimer’s disease.^58,59^ Full-length Tau is composed of an N-terminal acidic domain (residues 1-103), a proline-rich domain (residues 197-243), a microtubule-binding repeat domain comprising four tubulin-binding repeats (residues 244-368), which contains the aggregation-prone PHF6 and PHF6* motifs that drive Tau fibrillization, and a C-terminal domain (residues 369-441).^60^ Cryo-EM studies of Tau amyloid fibers derived from patient samples^61^ or self-assembled *in vitro*^62^ revealed a highly structured core formed from residues 306-378; the remaining 369 residues were not visible in cryo-EM reconstructions and are therefore presumably disordered and/or flexible.^61^ In contrast to amyloid fibrils from Tau, early-stage oligomers are not well studied,^63–65^ despite their cytotoxicity^66–68^ and suspected role in disease development by transmitting soluble nuclei of misfolded protein from cell to cell.^66,69,70^

The work presented here focuses on the characterization of transient mixed populations of early oligomers of Tau441(P301L) aggregated *in vitro*. First, we demonstrate the simultaneous assembly and insertion of PLY nanopores into planar phospholipid bilayers containing cholesterol. We characterize PLY nanopores and confirm their exceptionally stable and large inner diameter of 20 ± 3 nm; this size doubles the largest currently available biological nanopore diameter of poly(C9) pores.^19^ Second, we demonstrate the ability of PLY nanopores to estimate the volume and shape of individual folded proteins using a set of test proteins with molecular weights up to 103 kDa. Finally, we reveal the size and shape distribution of early oligomers of Tau as a function of incubation time, spanning the range from monomers to dodecamers of Tau441(P301L) with molecular weights ranging from 60 kDa to 660 kDa.

## Results and Discussion

### Self-assembly of PLY monomers to large nanopores and their characterization

To use PLY pores for nanopore experiments, we adapted previously reported protocols for PLY pore assembly on unilamellar, cholesterol-containing liposomes.^51,72–77^ Specifically, we pre-incubated purified PLY monomers (MW 53.7 kDa) with DTT and amphipol A8-35 and added this solution to the cis compartment of a recording chip with a planar lipid bilayer containing 30 mol% cholesterol directly on a recording chip (**Figure 1a**).

The solution of pre-incubated PLY and amphipol, in the absence of lipids, formed a few cylindrical PLY pore complexes in solution, which we used to characterize the inner diameter of the pore by transmission electron microscopy (TEM) (**Figure 1b** inset and **Figure S1**). Based on TEM images of these preparations, we estimate an average inner diameter of isolated PLY pores of 23.9 ± 0.8 nm (mean ± SD, *N* = 5), which is comparable to previously reported dimensions of the PLY pore obtained from cryo-EM studies of membrane-associated assemblies, reporting inner diameters of ∼ 26 nm^75^ and ∼ 31–50 nm depending on detergent conditions and conformational state of the pores.^76^

To assemble and insert the pore in a planar lipid bilayer, we added PLY-amphipol solution to the *cis* compartment of the recording chip at a final concentration of 3 µg mL^-1^ PLY, applied a potential of +100 mV to the *trans* side of the bilayer using two Ag/AgCl electrodes, and monitored pore incorporation *via* electrical recordings (**Figure 1a**). Typically, in a recording buffer with pH 7.5, PLY pores insert into the bilayers within 15 min, causing a sudden single-step increase in ionic current (**Figure 1b**). The magnitude of the current jump indicates an inner diameter of the membrane-reconstituted nanopores with a median value of 20.1 nm (19.7 ± 1.4 nm, mean ± SD, *N* = 50), corresponding to 35 ± 2 monomers of PLY (**Figure 1c, Supplementary Note 1**). Under weakly acidic or basic conditions (pH 6.5 or pH 8.0-8.5), we observed rare PLY pore insertions, which were characterized by a smaller and more varied pore diameter of 16 ± 6 nm (mean ± SD, *N* = 17) (**Supplementary Figure S2**).

For the characterization of the shape of natively folded proteins using nanopores, the electric field inside the nanopore must be uniform,^78^ and hence the nanopore must be cylindrical and remain so throughout the experiment. **Figure 1d** shows that the current−voltage relationship (I−V curve) of inserted PLY pores displayed Ohmic behaviour in a range of ionic strengths from 100 mM NaCl to 500 mM NaCl (**Supplementary Figure S3**), suggesting a cylindrical pore lumen. The schematic representation of the assembled PLY pore, shown in **Figure 1a**, and the TEM image in **Figure 1b** also indicate a cylindrical pore structure. **Figure 1d** shows that the pores were stable across a range of applied potentials from +100 mV to −100 mV. To assess the stability and associated noise characteristics of PLY nanopores, we compared the ionic current noise before and after pore insertion in a recording buffer containing 500 mM NaCl and 50 mM Tris−HCl, pH 7.5, based on power spectral densities (PSDs). The PSD graph shows only slightly increased noise in the low-frequency domain below ∼ 1 kHz (**Supplementary Figure S4**), suggesting fluctuations in the millisecond range within the inserted PLY assembly. Over time, the electrical noise of the PLY nanopore remained stable with excellent low-noise characteristics for resistive pulse recordings (**Figure 1b**). Most experiments ended approximately 30 min after pore insertion due to spontaneous rupture of the lipid bilayer; during this time, we typically recorded tens to hundreds of resistive pulses for protein characterization (**Figure 2**). Across all experiments, the membrane lifetime ranged from 12–30 min with an average of 22 ± 5 min (*N* = 23). We attributed a sudden increase in ionic current accompanied by amplifier saturation with a “current overload” to stochastic bilayer failure under conditions of applied voltage rather than to gradual pore destabilization; in some cases, we observed gradual stepwise increases in recorded current prior to rupture; these observations might have resulted from the insertion of additional PLY pores into the bilayer.

**Figure 2.**
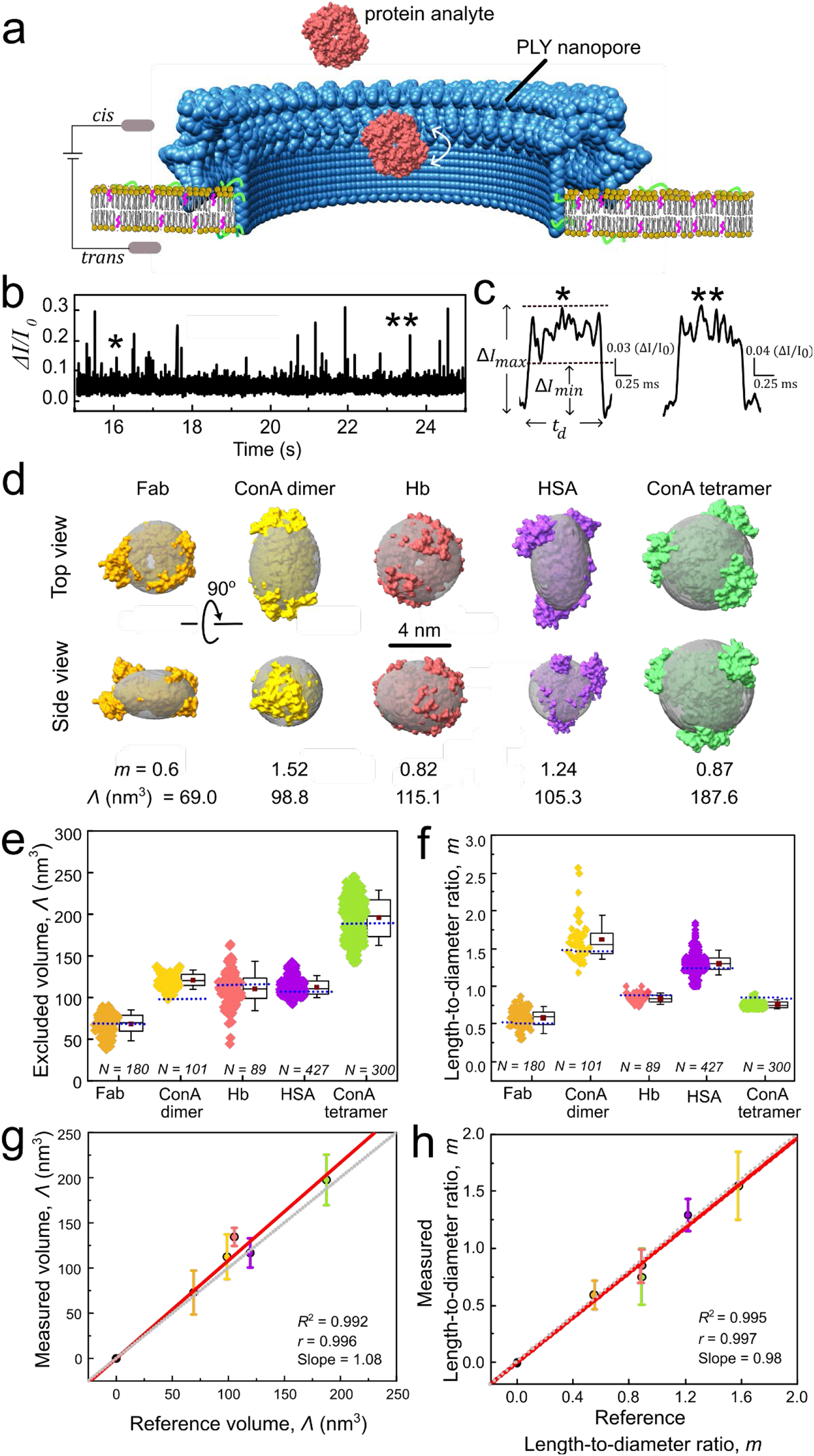
Estimation of the volume and shape (*i.e*., length-to-diameter ratio) of natively folded proteins based on resistive pulses recorded with PLY nanopores. **(a)** Schematic of the recording setup for single-molecule analysis using PLY pores embedded in a planar lipid bilayer. A cross-sectional view of the PLY nanopore is shown, along with a representative protein analyte inside the pore. (**b**) Baseline–corrected current recording in the presence of ConA dimers and tetramers, showing resistive pulses as upward spikes. Recording buffer, 500 mM NaCl, 10 mM Tris-HCl, pH 7.5, 0.02 µM amphipol, voltage = −100 mV applied to the bottom electrode in the *trans* compartment. The current recordings were collected with a 200 kHz sampling rate and filtered with a 20 kHz digital Gaussian low-pass filter for clarity. (**c**) Examples of two individual resistive pulses as marked in panel b, illustrating the maximum current blockage (Δ*I_max_*), minimum current blockage (Δ*I_min_*), and dwell time (*t_d_*). (**d**) Atomic structures of five test proteins and reference ellipsoids of revolution (in grey) used to estimate their expected volume *Λ* and shape *m*. Fab (PDB 7FAB) in orange, ConA dimer (PDB 1GKB) in yellow, Hb (PDB 1A3N) in red, HSA (PDB 1AO6) in purple, ConA tetramer (PDB 5CNA) in green. (**e**) Distribution of excluded volumes (*Λ*) and (**f**) length-to-diameter ratios (*m*) obtained from individual resistive pulses of five test proteins. Horizontal lines in the boxes represent median values of these distributions. Solid brown squares represent the experimental mean values. The box range represents the 25^th^ to 75^th^ percentile, and the whisker range displays the 10^th^ to 90^th^ percentile of these measured distributions. In contrast, blue horizontal dotted lines represent expected reference values from ellipsoid models of each protein. (**g, h**) Median values of the excluded volume (*Λ*), and the length-to-diameter ratio (*m*) determined from single event analyses of test proteins plotted against the expected reference values. Error bars show the 25^th^ to 75^th^ percentile. The grey dotted lines represent the ideal 1:1 agreement (slope = 1), and the red solid lines are the linear regressions performed imposing a zero intercept.

To assess the role of membrane composition in the probability of PLY pore insertion, we performed additional experiments inserting PLY pores into lipid bilayers composed of DiphyPC without cholesterol. Under these conditions, we observed only a low frequency of insertion: less than one insertion per hour, whereas in the presence of cholesterol, the frequency was consistently 4–6 insertions per hour. Additionally, in the absence of cholesterol, the average pore diameter was significantly smaller, 10.0 ± 5.2 nm (mean ± SD, *N* = 7, **Supplementary Figure S5**). Although recordings of PLY pores in cholesterol-free bilayers showed low noise and stable baselines, we found these conditions suboptimal due to the low insertion frequency and small pore diameter.

### Evaluation of PLY nanopores for estimating the volume and shape of analyte proteins

To test the performance of PLY nanopores for determining the volume and shape of single folded proteins, we used five test proteins: Fab fragment of polyclonal anti-biotin IgG (Fab, MW = 50 kDa), concanavalin A (ConA, MW dimer = 51.5 kDa, MW tetramer = 103 kDa, pI = 5.2), human hemoglobin (Hb, MW = 64.5 kDa, pI = 6.8) and human serum albumin (HSA, MW = 66.5 kDa, pI = 4.7). We added analyte proteins at a final concentration of ∼ 30 μg mL^-1^ to the *cis* side of the pore while applying a constant potential of −100 mV to the *trans* compartment of the lipid bilayer setup (**Figure 2a**). Resistive pulses in the current recordings indicated that all tested proteins were able to enter PLY nanopores (**Figure 2b**, **Supplementary Figure S6**). Analysis of the dwell time of resistive pulses *t*_d_ revealed that the most probable dwell times for Fab, ConA, Hb, and HSA were 180 μs, 295 μs, 315 μs, and 280 μs, respectively (**Supplementary Figure S7**). These residence times exceeded the diffusion time of a protein for the ∼ 10 nm length of a PLY pore by two orders of magnitude. These extended residence times allowed analyte proteins to sample various orientations within the pore and facilitated accurate estimation of protein volume and shape according to the theory developed by Golibersuch^79^ and others.^22,80^ Importantly, we did not observe resistive pulses in the absence of analyte proteins. Since the pH value of the recording buffer was above the pI values of most of the analyte proteins that we examined, these proteins were negatively charged during the experiments. For these proteins, we detected a higher frequency of resistive pulses when they migrated from a positively polarized *cis* compartment towards a negatively polarized *trans* compartment than from a negatively polarized *cis* compartment towards a positively polarized *trans* compartment. These results indicate that protein translocation through PLY pores was not dominated by electrophoretic motion of the negatively charged proteins but rather by electroosmotic flow through the net negatively charged lumen of PLY pores (**Figure 1a**). This conclusion is further supported by the voltage-dependence of the median translocation times shown in **Supplementary Figure S8**. We have previously made the same observation of EOF- dominated translocations with poly(C9) pores.^19^

To obtain quantitative estimates of protein volume and shape from resistive pulse recordings, both the pore diameter and length must be known. Since the measured open pore current depends on both parameters, it is important to determine the length of the pores as precisely as possible to update the diameter at any time during the recording based on the measured open pore current. Here, we took advantage of the fact that PLY nanopores represent a circular arrangement of identical protein subunits; therefore, the pore length is independent of the number of monomers in the assembly. Based on this assumption, we empirically determined the effective length of PLY nanopores using the five test proteins (**Figure 2d)**. To do so, we plotted the experimentally measured excluded volumes of Fab, ConA dimer and tetramer, Hb, and HSA as a function of their reference volumes while assuming various pore lengths ranging from 8 to 11 nm as a parameter in the analysis algorithm (**Supplementary Figure S9**). An effective pore length of 9.5 nm provided the best agreement with all five reference values, yielding a slope of 1.08 ± 0.06 and a regression coefficient of *r* = 0.996 (**Supplementary Figure S9**). Importantly, accurate determination of the volume and shape of analyte proteins did not require a constant pore diameter from experiment to experiment or even during the same experiment, because the known and constant effective length combined with the continuously measured open pore current between resistive pulses make it possible to update the pore diameter for the analysis of each resistive pulse.^19,78^

Using the effective pore length of 9.5 nm and the analysis algorithm published previously^19^ (**Supplementary Note 3**), we obtained median estimates for the excluded volume and length-to-diameter ratio for all test proteins (**Figures 2e-h, Supplementary Table S1)**. To evaluate the accuracy of the nanopore results, we compared them to the parameters of the reference ellipsoids of revolution for each protein. These reference values were calculated based on protein atomic structure as described in **Supplementary Note 4**. Briefly, each protein surface was estimated by the ellipsoid with axes A, B, B, *m* = A/B with *m* < 1 corresponding to an oblate and *m* > 1 to a prolate shape.^27^

We observed excellent agreement between the experimental and reference parameters (**Figures 2e-h**). The reference parameters are deterministic values calculated directly from the corresponding PDB structures of each protein and therefore represent single expected values rather than distributions. Additionally, we tested alternative approaches to calculating reference values. A different algorithm for ellipsoid approximation from atomic structures^81^ provided similar results, whereas estimation of reference volumes based solely on molecular weight and the assumption of a perfectly spherical protein shape resulted in a systematic underestimation of the excluded volume by approximately minus 30% (**Supplementary Table S1**). In this context, the PLY pore with a pore diameter of ∼ 20 nm extends the regime of large biological nanopores from PlyAB pores with a diameter of ∼ 5.5 nm and the membrane attack complex (polyC9 with a pore diameter of ∼ 10 nm); these biological nanopores are distinct from small nanopores such as α-hemolysin, MspA, or aerolysin.

### Molecular weight characterization and kinetic modelling of Tau oligomerization

To demonstrate the potential of PLY nanopores for the characterization of clinically relevant protein aggregates, we investigated their ability to characterize Tau oligomers on a single particle level. We used Tau441(P301L), the longest, brain-specific isoform of Tau in humans, comprising 441 amino acids (**Figure 3a**). Although its atomic structure has not been experimentally determined, recently reported discrete molecular dynamics simulations guided by short-range crosslinking constraints provide a reference structure in which Tau441 monomers adopt a compact conformation, characterized by transient, dynamic secondary structure elements (**Figure 3b**).^82^ To investigate the kinetics of aggregation of Tau441(P301L), we added heparin at a Tau : heparin molar ratio of 2:1 to induce aggregation at 37°C;^83^ heparin is a well-established promoter of Tau fibrillation.^84^ **Figure 3c**, shows the abundance of all Tau441(P301L) species from the monomer to the dodecamers at various time points of aggregation as determined by resistive pulse analysis with PLY nanopores (see **Supplementary Figure S10** for the corresponding current recordings and **Supplementary Figure S11** for typical current modulation within single resistive pulses).

**Figure 3.**
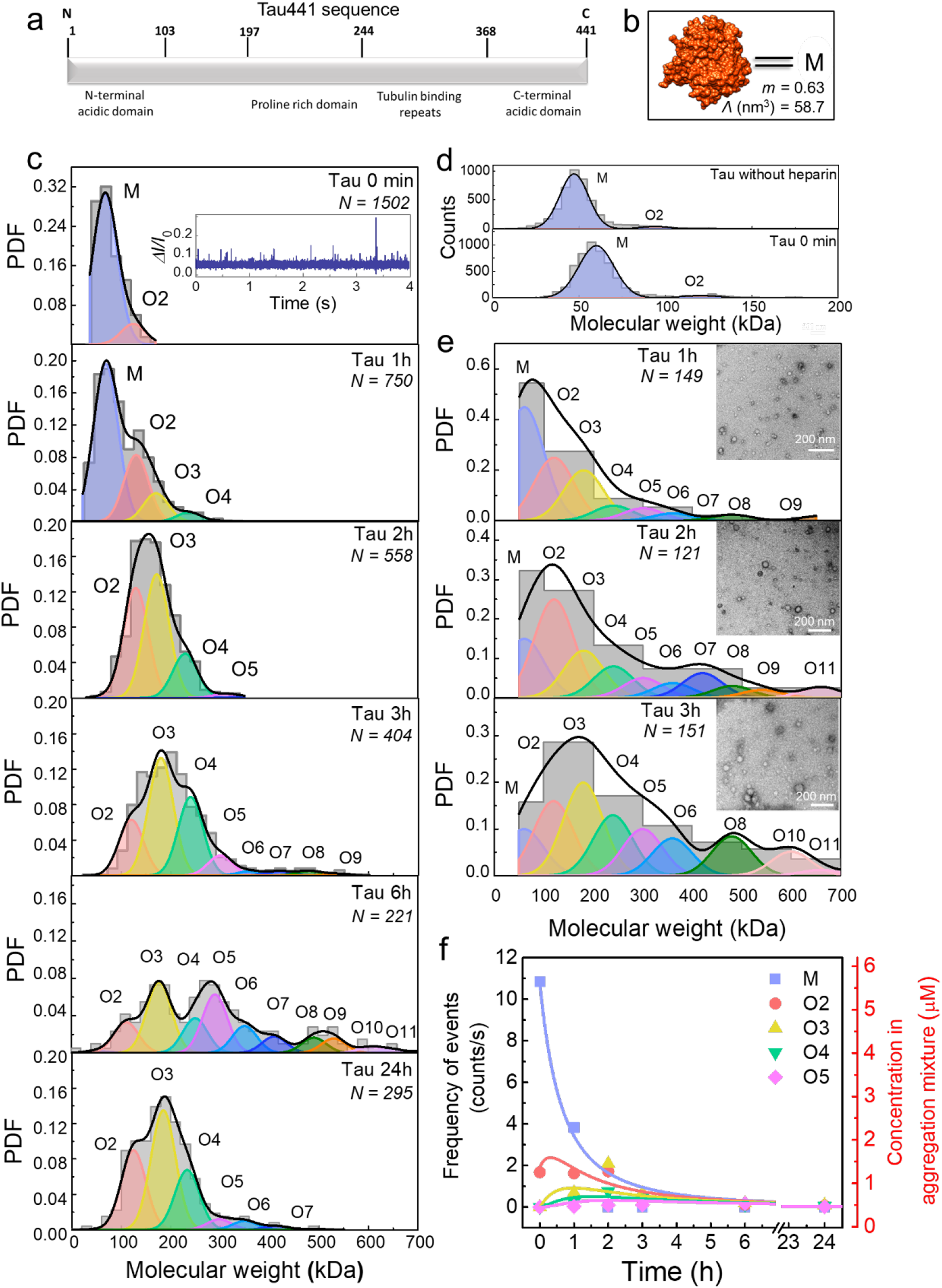
Characterization of individual unlabeled Tau oligomers with PLY nanopores. (**a**) Illustration of the amino acid sequence of Tau441(P301L) showing the positions of the acidic domains (N- and C-terminal), and tubulin binding repeats. (**b**) Surface representation of conformational ensemble of Tau441(P301L) monomer (PDB 8ZZX), as determined by discrete molecular dynamics (CL-DMD) simulations.^82^ (**c**) Time-dependent molecular weight distributions of subpopulations of Tau oligomers from monomers to dodecamers during aggregation as determined from resistive pulses with PLY nanopores. Molecular weights were determined from measured excluded volumes (*Λ*, nm^3^)^85^ (see **Supplementary Note 5**). Tau oligomer sub-populations are labeled from monomers (M) to oligomers containing eleven monomers (O11), corresponding to molecular weights from 60 kDa (M) to 660 kDa (O11). The inset shows baseline–corrected current recordings with resistive pulses as upward spikes at the 0-minute time point during Tau aggregation. Data were obtained from at least three independent PLY pores at each time point. The recording buffer contained 500 mM NaCl and 50 mM Tris-HCl, pH 7.5. (**d**) Molecular weight distributions of Tau without heparin and in the presence of heparin (Tau 0 min) as determined by mass photometry in 50 mM Tris-HCl recording buffer containing 500 mM NaCl with a pH of 7.5. (**e**) Time-dependent molecular weight distributions of Tau oligomers determined by TEM with corresponding TEM micrographs of Tau oligomers (insets). The molecular weight of oligomers was estimated^85^ using the area-equivalent circular diameter calculated from the major and minor axes of the 2D particle projections (see **Supplementary Note 5** and **Supplementary Figures S13-S15**). (**f**) Time evolution of the abundance of various subpopulations of Tau oligomers in solution. Frequency of resistive pulses (left y-axis) and concentration of Tau (right y-axis) as determined from current recordings in the presence of various Tau oligomers using PLY nanopores. The continuous lines are fitted to the experimentally determined frequencies using a colloidal aggregation model described in **Supplementary Note 8**.

First, we analysed the volume of Tau441(P301L) monomers at the 0 min time point. The median excluded volume determined with PLY nanopores was 70 ± 9 nm^3^, which was significantly larger than the reference volume of 58.7 nm^3^. Conversion of the experimentally measured volume to molecular weight (**Supplementary Note 5**)^85^ yielded an apparent molecular weight of 60.0 ± 7.1 kDa (**Figure 3c**), exceeding the expected reference value of 45 kDa by approximately 15 kDa, which corresponds to the average molecular weight of the heparin molecules used in our preparation. The nanopore data therefore suggest that Tau441 (P301L) monomers form a relatively stable 1:1 complex with heparin, as reported previously.^84^

To validate this interpretation, we performed mass photometry measurements of Tau441(P301L) monomers in the presence and absence of heparin in an amphipol-free recording buffer. Mass photometry yielded distinct mass estimates of 45 ± 11 kDa for Tau alone and 60 ± 10 kDa for Tau in the presence of heparin (**Figure 3d**), supporting a 1:1 stoichiometry of the Tau-heparin complex. Additionally, at the 0-minute time point, both PLY nanopore and mass photometry measurements revealed a small subpopulation of dimers (O2), with no higher-order oligomers detected (**Figure 3c-d**). Such sample heterogeneity is consistent with previously reported data.^86^

We next analysed the molecular weight distributions of Tau oligomers after incubation at 37°C in the presence of heparin over time (**Figure 3c).** With increasing incubation time, monomers disappeared while progressively higher-order oligomers with molecular weights up to ∼660 kDa emerged. To assess which oligomeric subpopulations contributed to these distributions, we fitted the molecular weight distributions using multi-component Gaussian fitting with the following constraints: the peak widths were fitted to one value shared across all components, and peak positions were restricted to *N* ∗ 60 *kDa* ± 10 *kDa*, where *N* denotes the aggregation number (*i.e*., the number of Tau441(P301L) monomers in each oligomer). This approach enabled consistent assignment of monomers, dimers, and higher-order species across time points. The oligomer distributions reported here reflect the heparin-induced Tau aggregation pathway under the experimental conditions used in this study.^59^

In parallel with the progression of aggregation toward larger species, the frequency of resistive pulses decreased with incubation time, leading to fewer oligomer particles detected at later stages of aggregation (**Figure 3c**). This reduction reflects, on the one hand, a decrease in the total oligomer number, as small oligomers are increasingly sequestered into larger oligomers during later stages of aggregation, and, on the other hand, the formation of large oligomers (> O11) that cannot enter the pore. The nanopore measurements primarily report on soluble, pore-accessible Tau oligomers throughout the aggregation process. Larger insoluble aggregates and mature fibrillar species are likely underrepresented or absent in the nanopore measurements.

Complementary techniques for sampling early oligomeric Tau populations, including TEM imaging and mass photometry, agree well with nanopore data (**Figure 3e, Supplementary Figure S13**). Visualization by TEM revealed a heterogeneous mixture of globular Tau oligomers (**Figure 3e** insets and **Supplementary Figure S14**). The diameters of these particles were measured using Feret’s diameter in ImageJ and converted into estimated protein molecular weights (**Figure 3e**, **Supplementary Figure S15, Supplementary Note 5**). Approximately 90% of the oligomers detected by TEM had molecular weights below 700 kDa and were accessible to nanopore measurements with Ply pores. The full molecular-weight distribution determined by TEM included only a minor fraction (∼10%) of higher-order oligomers (**Supplementary Figure S16**).

Mass photometry measurements also revealed the formation of oligomeric species with molecular weights up to 700 kDa at extended incubation times (**Supplementary Figure S13**). Notably, the oligomer distributions determined by mass photometry showed more pronounced monomer and dimer peaks. This observation may, in part, reflect a bias due to the speed of diffusion of oligomers of different sizes to the detector surface during the short acquisition time (1 min). The overall trend in oligomer formation, however, is consistent with the nanopore data.

The toxicity of Tau oligomers depends on their size^87^ and, possibly, their shape.^88^ Several groups reported that small soluble oligomers of Tau in the range from dimers to decamers are among the most toxic aggregates.^88–90^ To analyze the kinetics of the formation of low-n Tau oligomers based on the nanopore data, we determined the event frequency of each oligomer species at various time points. Specifically, we quantified the total number of resistive pulses for each oligomeric peak in **Figure 3c** and normalized them to the total duration of the recordings that revealed these pulses. To describe the early stage of aggregation of low-n oligomers, we fitted the event frequencies of monomers, dimers, trimers, tetramers, and pentamers at different time points in **Figure 3f** with a kinetic model that describes aggregation as a set of irreversible pairwise collisions between all possible species present in solution (**Supplementary Note 8**). In this model, the rates of individual pairwise collisions depend solely on the sizes and diffusion coefficients of the oligomers, while a single energy barrier (*E*_a_) serves as the only fitting parameter, accounting for steric and electrostatic effects that may slow aggregation relative to a diffusion-limited process. To avoid overfitting, we assumed *E*_a_ to be identical for all colliding oligomers and obtained the best fit at *E*_a_ = 10.5 *k*_B_*T* .A similar analysis of the time-dependent oligomer abundances in the corresponding TEM data using the same kinetic model yielded a comparable activation barrier of *E*_a_ = 10.7 *k*_B_*T* , in close agreement with the nanopore-derived value and supporting the consistency of the time-resolved aggregation analysis (**Supporting Figure S17**, see **Supplementary Note 8** for details). This value is physically reasonable and indicates slow aggregation kinetics that are not dominated by diffusion. The kinetic model provides a descriptive representation of the early evolution of the detectable oligomer population and is not intended to identify aggregation nuclei or mechanistically distinguish on-pathway from off-pathway oligomeric species. While this model is likely oversimplified, it describes the experimental data on the formation of the smallest (< 10-mer) – often considered most toxic – oligomers^88^ fairly well (**Figure 3f**). **Figure 3f** also reveals the concentrations of low-n oligomers from dimers to pentamers as determined from the frequencies of resistive pulses and their corresponding amplitudes of current blockade (**Supplementary Note 9**). According to Arosio, Knowles, and Linse, quantifying the concentrations of subpopulations of all different oligomers in situ is considered one of the major unmet challenges in protein amyloid aggregation studies.^87^ The quantification of the concentrations of low-n oligomers of different size (**Figure 3**) and shape (**Figure 4**) presented here addresses this unmet need and may enable correlation studies between oligomer size, shape, and toxicity, as reported previously by Prangkio *et. al.* for low-n oligomers of amyloid- β.^29^ These measurements reflect the subset of soluble, pore-accessible oligomeric species detectable by the nanopore and therefore do not represent the full spectrum of Tau aggregation states.

**Figure 4.**
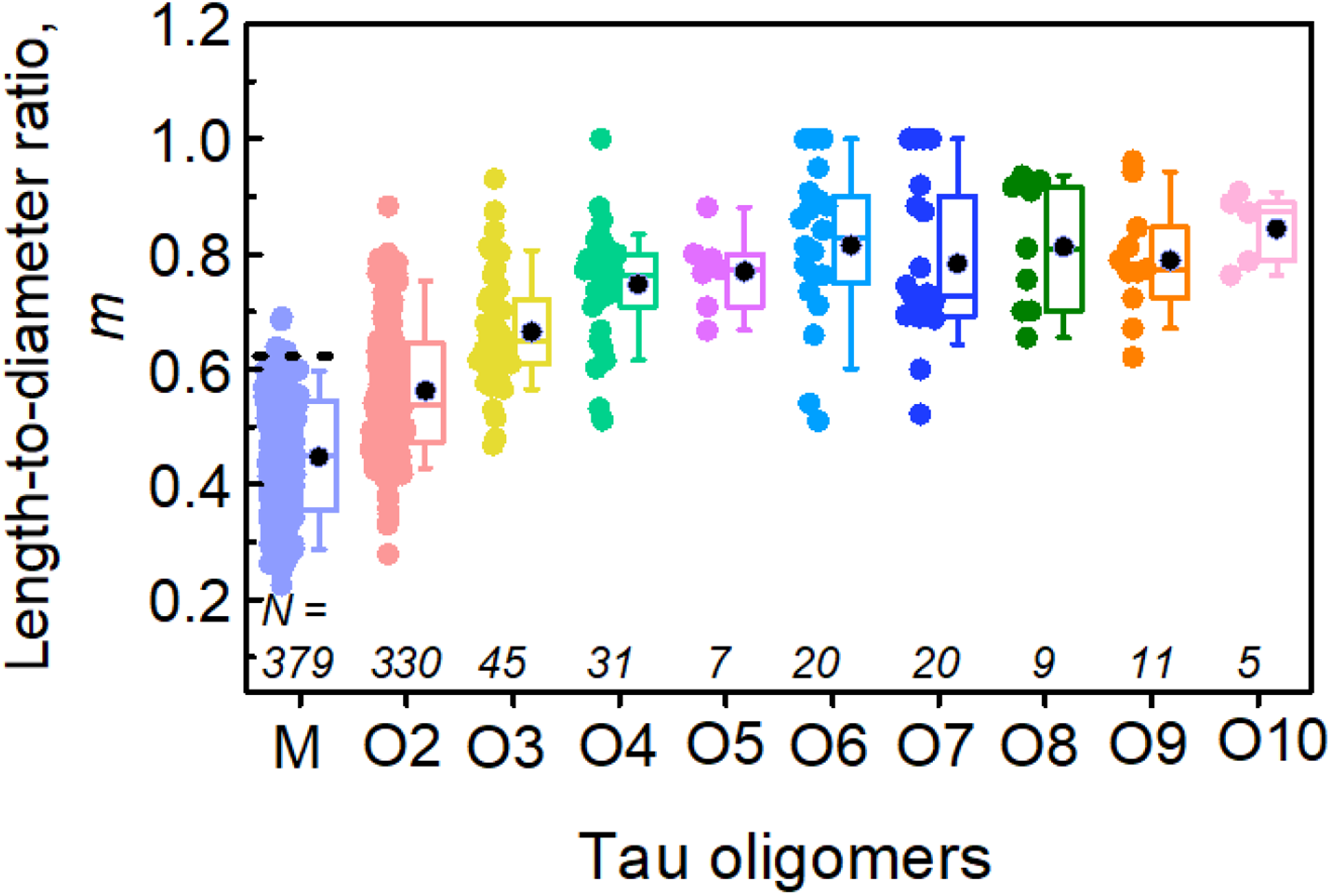
Single-molecule shape estimation of Tau oligomers in solution. Combined values of length-to-diameter ratios (*m*) determined from resistive pulses of Tau oligomers during aggregation at varying time points (0 to 24h) using PLY nanopores and a recording buffer containing 500 mM NaCl, Tris-HCl, pH 7.5. The black horizontal dotted line represents the expected reference value of Tau monomer (M) from the ellipsoid model.

### Characterization of the shape of individual Tau oligomers

Finally, we estimated the length-to-diameter ratio, *m,* of each oligomeric subpopulation (**Figure 4**), assuming *m < 1,* corresponding to an oblate shape as observed by TEM (**Figure 3e**, insets).^22,27^ For analysis, we pooled data across all time points and selected the resistive pulses corresponding to *N* ∗ 60 ± 8.2 *kDa*, where *N* is the aggregation number of the oligomer, to exclude ambiguous events that reveal molecular weights when oligomer subpopulations overlap. Additionally, only long resistive pulses with dwell times *t*_d_ > 450 µs were included in the shape analysis to acquire resistive pulse modulations from a representative sample of particle orientations (**Supplementary Figure S18**).^22^

**Figure 4** shows the distribution of the length-to-diameter ratio *m* across species ranging from Tau monomers (M) to the highest-order oligomers detected (O10). A value of *m* ≈ 1 corresponds to a sphere, whereas lower values indicate increasingly oblate structures. The nanopore measurements revealed a progressive increase in *m* from ∼0.45 for monomers to ∼0.85 for O10 oligomers, indicating that larger assemblies adopt more compact geometries. These results demonstrate that the soluble, pore-accessible Tau oligomers detectable by the nanopore undergo a gradual structural transition from predominantly oblate-shaped Tau441(P301L) monomers, dimers, and trimers toward more spherical tetramers to decamers. Within this size range, which has previously been implicated in neurotoxicity,^91–94^ the nanopore experiments carried out here do not observe a transition to elongated prolate shapes during the course of the experiment.^91–94^ The ability to estimate the size and, in particular, the shape of individual oligomeric species inside the solution mixture of a heterogeneous sample of amyloidogenic proteins is a unique analytical capability of PLY nanopores.

## Conclusion

In this work, we introduce PLY nanopores for the label-free single-particle analysis of large, folded proteins and their oligomeric assemblies in heterogeneous solution mixtures. We demonstrated stable assembly and insertion of PLY pores into cholesterol-containing planar lipid bilayers and confirmed their exceptionally large inner diameter of 20.2 ± 3.0 nm and effective pore length of 9.5 nm, enabling sensitive detection of macromolecular species well beyond the size range accessible to previously published biological nanopores. Using PLY nanopores for resistive pulse analysis of reference proteins with known atomic structures, we demonstrate the quantification of excluded volume, molecular weight, and shape of single proteins and protein aggregates with exceptional accuracy. One of the most important features of PLY pores is that the label-free analysis of single protein particles that they afford is calibration-free, and that the results from experiments on different days and from different operators are directly comparable. The reason for this benefit is that the effective length of PLY nanopores is always constant, and that the pore diameter is readily monitored and updated throughout the experiment. With knowledge of the effective pore length, pore diameter, and electrolyte conductivity, the volume and shape of analyte proteins are revealed by the magnitude and modulations of their resistive pulses. Importantly, the determination of protein volume and shape does not require calibration against a reference protein, as the length of PLY pores is constant and known, the shape of the pore can be approximated by a cylinder, the conductivity of the recording electrolyte is known and constant, and the continuously measured open-pore conductance reveals the pore diameter at all times during the experiment. This characteristic is a dramatic advantage compared to solid-state nanopores, since these pores must be calibrated before each experiment, as slow etching of the SiN_x_ windows that contain the nanopores increases their diameter over time, and as the shape of the nanopore lumen may deviate significantly from a cylindrical shape that is ideal for accurate estimation of protein shape.^28^

Applying PLY nanopores to Tau aggregation, we detected monomers, dimers, and progressively higher-order oligomers over time, reaching molecular weights of up to ∼660 kDa corresponding to dodecamers. At the earliest time point, the apparent molecular weight of Tau was consistent with the formation of a 1:1 Tau–heparin complex. At later time points, we identified the relative abundance and absolute concentrations of low-n oligomers of Tau, which are associated with cytotoxicity. A kinetic model of the very first stages of aggregation based on irreversible pairwise collisions with a single activation energy barrier *E*_a_ = 10.5 *k*_B_*T* suggested that Tau aggregation was not diffusion-limited. In addition, PLY nanopores have the beneficial characteristic that the most probable residence time of all tested translocating proteins was in the range of hundreds of microseconds (see **Supplementary Figure S7**). This characteristic is fortuitous as it provides a large fraction of long resistive pulses for quantitative shape analysis of individual oligomeric species within a heterogeneous sample. Together, these results demonstrate that PLY nanopores enable label-free, quantitative characterization of protein oligomers in the clinically relevant size range at the single-particle level and provide a tool with unique, enabling capabilities for studying the early stages of protein aggregation.

## Methods

### Purification of Pneumolysin (PLY) monomers

We expressed PLY monomers in E. coli and purified them as previously described.^74,76,95^ Briefly, we amplified the PLY gene from *Streptococcus pneumoniae* D39 (Clontech, Saint-Germainen-Laye, France), cut the PCR products and the pET28a vector (Novagen, Madison, USA) with restriction enzymes (BamHI, XhoI, Fisher Scientific, Wohlen, Switzerland), and ligated them using T4-DNA ligase (Promega, Switzerland). We transformed the vector into BL21 (DE3) pLysS competent cells (Promega, Switzerland) and confirmed positive clones by sequencing. We induced protein expression with 1 mM IPTG (Sigma-Aldrich) at an OD_500_ of 0.5–0.7 in the bacterial culture. We harvested bacterial cells after 4h incubation at 37 °C and resuspended in a lysis buffer containing Lysozyme (Sigma-Aldrich) and DNAse I (Sigma-Aldrich). To lyse the cells, we sonicated them on ice in 10 × 20 s steps with 15 s breaks (BANDELIN sonoplus uw 2070, BANDELIN Electronic GmbH & Co. KG). We centrifugated the lysate at 70.000 ×*g* for 1 h at 4 °C and purified the recombinant proteins from the supernatant with 1 mL Ni-NTA columns (MACHEREY-NAGEL, Oensingen). In the final step, we dialyzed the sample against 50 mM Tris-HCl buffer (pH 7.0) to obtain a final concentration of 1 mg mL^-1^.

### Activation of PLY monomers and treatment with amphipol

We activated purified PLY monomers (1 mg mL^-1^, 5 µL) by incubating them with an equal volume (5 µL) of 50 mM Tris−HCl buffer (pH 7.5, Thermo Scientific) supplemented with 15 mM 1,4-dithiothreitol (DTT, Sigma-Aldrich 10197777001) for 10 min at 37 ⁰C. We then incubated the activated PLY monomers with amphipol A8-35 (Anatrace), maintaining a PLY:amphipol molar ratio of 1:2 in 50 mM Tris−HCl (pH 7.5) overnight at 37 ⁰C. The final concentration of PLY monomers was 0.25 mg mL^-1^.

### Planar lipid bilayer formation and characterization

We prepared a lipid mixture containing 70 mol% 1,2-diphytanoyl-sn-glycero-3-phosphocholine (DiphyPC, Sigma-Aldrich/Merk 850356P) and 30 mol% cholesterol (Fisher Scientific 11493310) in octane (Sigma-Aldrich/Merk 296988) with a final concentration of 15 mg mL^-1^ DiphyPC. For planar lipid bilayer current measurements, we used an integrated chip-based, four-channel parallel bilayer recording setup (Orbit Mini, Nanion Technologies) with multielectrode cavity-array chips (MECA chips, Ionera)^96^ and EDR3 software (Elements). The MECA chips, bearing four channels with a cavity diameter of 150 µm, supported lipid bilayers that we formed following a previously described technique.^96^ Briefly, we added 150 µL of a recording buffer (500 mM NaCl, 50 mM Tris−HCl pH 7.5, 0.02 mg mL^-1^ amphipol A8-35) to the *cis* compartment of the MECA chip and wetted all four channels by applying gentle pressure with a syringe plunger. Lipid bilayers were then painted manually by applying 0.3–0.4 µL of the lipid mixture near the cavities on the chip, as described below. A pipet tip was dipped into the lipid mixture solution without pipetting. The tip was then carefully dried with a tissue paper, leaving only a trace amount of lipids on the tip surface. Next, it was placed in the vicinity of a cavity on the chip and pipetted once to make an air bubble on the cavity. Once the lipid bilayer was formed, we aspirated the air bubble from the cavity. We confirmed the quality of the lipid bilayers by verifying that the baseline current stayed in the range of 0 ± 0.3 nA while applying ± 100 mV potential and the electrical capacitance in the range 30 ± 5 nF. The recording software EDR3 automatically estimated the membrane capacitance by analyzing the current response to an applied triangular potential difference. We allowed the bilayers to stabilize for 5 min. During this time, we tested the stability of the bilayers by applying transmembrane voltages of ±100 mV for 1 min and confirming the expected noise level and the absence of leak currents.

### In situ PLY nanopore assembly in a planar lipid bilayer

We added 2.5 μL of 0.2 mg mL^-1^ PLY-amphipol solution to the *cis* compartment of the MECA chip. We maintained an applied potential of +100 mV until a sudden single-step current jump was observed, indicating successful assembly or insertion of PLY nanopore in the lipid bilayer. During a typical experiment, a nanopore formed within 10–15 min after the addition of the PLY-amphipol solution. The inner diameter of the pore and the corresponding number of PLY monomers were calculated from the pore conductance measured as a difference between baseline and open pore current. We estimated the pore diameter using the equation proposed by Cruickshank *et al*,^97^ which assumes the pore to be perfectly cylindrical and accounts for both the resistance of the nanopore and the access resistances:

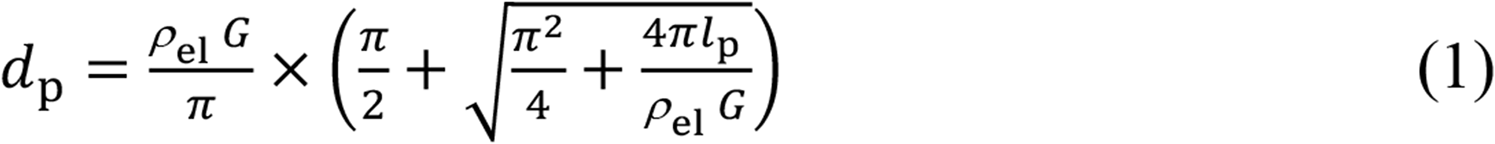

In equation 1, *d*_p_ [m] is the inner diameter of the nanopore, *l*_p_ [m] is the pore length, *ρ*_el_ [Ω·m] is the electrical resistivity of the electrolyte buffer and *G* [S] is the conductance of the pore. We used the channel length *l*_p_ = 9.5·10^-9^ m, determined as the effective pore length of PLY pores (**Supplementary Figure S9**). We then calculated the number of PLY monomers composing the nanopore using a previously published geometric model.^19^ We conducted all electrical measurements at a sampling rate of 200 kHz and a current range of 20 nA selected via the EDR3 software.

### In vitro aggregation of Tau protein

We carried out *in vitro* aggregation of Tau441 following a protocol reported by Crespo *et. al*.^98^ Briefly, we resuspended dry Tau441 powder (P301L mutant, rPeptide) to a concentration of 0.33 mg mL^-1^ in aggregation buffer containing PBS (pH 7.1) and 0.5 mM Tris (2-carboxyethyl) phosphine (TCEP, Sigma-Aldrich). We then added heparin (Sigma-Aldrich, H3393-100KU, 14 kDa) to the Tau solution in a 1:2 molar ratio and incubated the mixture at 37 °C and 500 rpm in a ThermoMixer (Eppendorf) for aggregation. We collected samples at different times, flash-froze 10 µL aliquots in liquid nitrogen, and stored them at −80 °C for future experiments.

### Resistive pulse sensing experiments with protein analytes

After inserting a PLY nanopore into a planar lipid bilayer, we performed a stepwise potential sweep from −100 mV to +100 mV to confirm pore stability and determine the offset current at 0 mV applied potential. Subsequently, we added 5 µL of the analyte protein solution to the *cis* compartment of the chip and the final protein concentration is mentioned in the section “Handling of analyte proteins” below. To promote translocation of the analyte proteins through the nanopore, we applied a constant ±100 mV potential across the bilayer. Every 5 min, we repeated the potential sweep to verify nanopore stability and to determine the possible changes in offset potential. If an offset potential was present, we corrected for it with respect to the recorded currents.

### Excluded volume analysis from resistive pulse measurements

We determined the excluded volume of individual protein analytes from the measured current blockade Δ*I* generated by resistive pulses through PLY nanopores. To this end, we first determined the radius of the nanopore from the recorded open pore current (*I*_0_) at a known applied potential *V* (i.e., from the open pore conductance), using equation 2, which accounts for both the resistance of the nanopore and its access resistance at the pore entrance and exit:

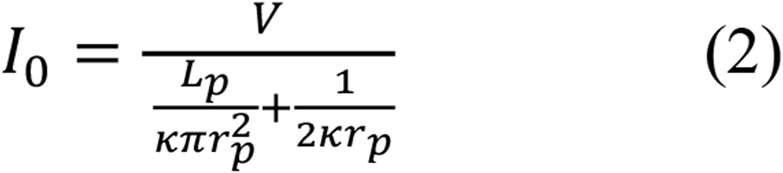

In equation 2, *V* (V) is the applied potential, *L_p_* (m) is the effective pore length, *r_p_* (m) is the pore radius, and κ (S m^-1^) is the conductivity of the electrolyte. We determined the normalized current blockade (Δ*I*/*I*_0_) for each resistive pulse and calculated the excluded volume (Λ) of the analytes according to equation 3:

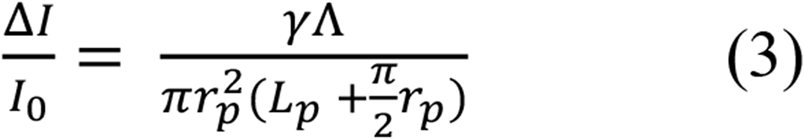

In equation 3, γ is the electrical shape factor and *Λ* (nm^3^) is the excluded volume of the analyte. We used an effective pore length of *L_p_* = 9.5 nm, which we determined from calibration measurements using proteins with known molecular volumes (**Supplementary Figure S9**), for all calculations. We continuously monitored the open pore conductance during each experiment, thereby accounting for pore-to-pore variations in diameter between experiments and potential changes in pore dimensions during recordings. This approach made it possible to determine excluded volumes on an event-by-event basis from individual resistive pulses.

### Handling of analyte proteins

We used a solution Fab protein (Fab fragment of polyclonal anti-Biotin IgG from Goat, Thermo Scientific, cat. # 800-101-098) in nanopore experiments at a final concentration of 0.66 μM in the *cis* compartment of the MECA chip. Human serum albumin (HSA, Sigma-Aldrich, A1653), Haemoglobin (Hb, Sigma-Aldrich, H7379), and Concanavalin A (ConA, Sigma-Aldrich, C2010) were resuspended in PBS buffer (ROTI Cell, 9143), pH 7.5, to prepare the stock protein solutions. The final molar concentrations of analyte proteins in the cis compartment were 0.50 μM for HSA, 0.51 μM for Hb, and 0.72 μM ConA (calculated per monomer equivalent). In the case of the Tau441(P301L) protein, we added 2-3 μL of the aggregation mixture to a volume of 150 – 200 μL of recording buffer in the cis compartment.

### Analysis of nanopore data

We filtered the acquired data with a Gaussian low-pass filter at a cut-off frequency of 20 kHz and performed a threshold search (5 × the standard deviation of the baseline current) for the detection of resistive pulses.

### Transmission electron microscopy (TEM)

We used an oxygen plasma cleaner (Zepto Rie, Diener electronic) to glow-discharge carbon-coated 300-mesh copper grids (Electron Microscopy Sciences) for 5 s, ensuring that the grid was clean and negatively charged. We applied a 5 μL drop of a 2 µg mL^-1^ PLY- amphipol solution to the freshly glow-discharged grid for 1 min and washed with milli-Q water to remove unbound protein. We further incubated the grid with 2 μL of 2% v/v aqueous uranyl acetate solution for 5 min, blotted off excess stain with a filter paper, washed the grid with 10 μL milli-Q water, and dried under ambient conditions for 30 min. We performed TEM imaging using a Tecnai G2 electron microscope (FEI) at an operating voltage of 120 kV. We used ImageJ^99^ to measure the inner diameter of the PLY nanopore and Tau samples. See **Supplementary Note 6** and **Supplementary Figure S16** for Tau oligomer molecular weight distribution analysis.

### Mass Photometry (MP) analysis

We acquired all the mass photometry data using a TwoMP mass photometer (Refeyn Ltd., Oxford, UK), similar to that described previously by Paul *et al*. with slight modifications.^86^ Briefly, we cleaned microscope coverslips (24 × 50 mm Thorlabs, cat. no. CG15KH1) using isopropanol followed by three washes with pure water. We used PDMS CultureWell gaskets (cat. no. GBL103250) to define compartments for single-oligomer mass analysis. We diluted Tau solutions to a concentration of ∼280 ng mL^-1^ in 50 mM Tris-HCl recording buffer containing 500 mM NaCl with a pH of 7.5 immediately prior to mass photometry measurements. For each mass photometry acquisition, 20 μL of diluted protein was introduced into the chamber, and following autofocus stabilization, movies of 60 s duration were recorded. Each sample was measured at least three times independently. Mass photometry data acquisition was performed using AcquireMP software (Refeyn Ltd., v2.4.1) with large view acquisition setup. Contrast-to-mass calibration was performed using MassFerence P1 Calibrant (Refeyn Ltd., MP-CON-41033). The histograms (bin size 5 kDa) were fitted with Gaussian distributions in the range of 25–700 kDa.

## AUTHOR INFORMATION

Notes

The authors declare the following competing interests: Anasua Mukhopadhyay, Wachara Chanakul, Yu-Noel Larpin, Saurabh Awasthi, Alessandro Ianiro, and Michael Mayer have filed a patent (applicant: Adolphe Merkle Institute, University of Fribourg) with the title of PLY NANOPORES (PCT/EP2024/084743). We declare no further competing financial interests.

## ACKNOWLEDGMENTS

A. M. acknowledges financial support by the Swiss National Science Foundation (SNSF) “SPARK” funding (CRSK-2_221078), support from the SNSF through the National Center of Competence in Research (NCCR) Bio-Inspired Materials, Novartis Foundation for Medical-Biological Research, as well as the Adolphe Merkle Institute. W. C. acknowledges a Swiss Government Excellence Scholarship (2020.0624) to complete his Ph.D. at the Adolphe Merkle Institute, University of Fribourg. S. A. acknowledges financial support from the Department of Biotechnology, India (ref. no: BT/RLF/Re-Entry/40/2021). A. I. acknowledges support from the SNSF through the NCCR Bio-Inspired Materials. M. M. acknowledges financial support from the SNSF (Grant number: 200020_197239), the SNSF NCCR Bio-Inspired Materials, and the Adolphe Merkle Foundation, Switzerland.

## Data availability

The data presented is available in the Zenodo repository: https://doi.org/10.5281/zenodo.22035618. Source data are provided with this paper.

## References

1. Bayley, H. & Martin, C. R. Resistive-Pulse Sensing From Microbes to Molecules. Chem. Rev. 100, 2575–2594 (2000).

2. Robertson, J. W. F. & Reiner, J. E. The Utility of Nanopore Technology for Protein and Peptide Sensing. Proteomics 18, 1800026 (2018).

3. Restrepo-Pérez, L., Joo, C. & Dekker, C. Paving the way to single-molecule protein sequencing. Nat. Nanotechnol. 13, 786–796 (2018).

4. Wanunu, M., Morrison, W., Rabin, Y., Grosberg, A. Y. & Meller, A. Electrostatic focusing of unlabelled DNA into nanoscale pores using a salt gradient. Nat. Nanotechnol. 5, 160–165 (2010).

5. Motone, K. et al. Multi-pass, single-molecule nanopore reading of long protein strands. Nature 633, 662–669 (2024).

6. Wanunu, M. et al. Rapid electronic detection of probe-specific microRNAs using thin nanopore sensors. Nat. Nanotechnol. 5, 807–814 (2010).

7. Keyser, U. F. et al. Direct force measurements on DNA in a solid-state nanopore. Nat. Phys. 2, 473–477 (2006).

8. Plesa, C. et al. Fast Translocation of Proteins through Solid State Nanopores. Nano Lett. 13, 658–663 (2013).

9. Hu, R. et al. Intrinsic and membrane-facilitated α-synuclein oligomerization revealed by label-free detection through solid-state nanopores. Sci. Rep. 6, 20776 (2016).

10. Baaken, G. et al. High-Resolution Size-Discrimination of Single Nonionic Synthetic Polymers with a Highly Charged Biological Nanopore. ACS Nano 9, 6443–6449 (2015).

11. Hu, Z., Huo, M., Ying, Y. & Long, Y. Biological Nanopore Approach for Single-Molecule Protein Sequencing. Angewandte Chemie International Edition 60, 14738–14749 (2021).

12. Mayer, S. F., Cao, C. & Dal Peraro, M. Biological nanopores for single-molecule sensing. iScience 25, 104145 (2022).

13. Maglia, G., Restrepo, M. R., Mikhailova, E. & Bayley, H. Enhanced translocation of single DNA molecules through α-hemolysin nanopores by manipulation of internal charge. Proceedings of the National Academy of Sciences 105, 19720–19725 (2008).

14. Ying, Y.-L. et al. A Time-Resolved Single-Molecular Train Based on Aerolysin Nanopore. Chem 4, 1893–1901 (2018).

15. Kasianowicz, J. J., Brandin, E., Branton, D. & Deamer, D. W. Characterization of individual polynucleotide molecules using a membrane channel. Proceedings of the National Academy of Sciences 93, 13770–13773 (1996).

16. Aksimentiev, A. & Schulten, K. Imaging α-Hemolysin with Molecular Dynamics: Ionic Conductance, Osmotic Permeability, and the Electrostatic Potential Map. Biophys. J. 88, 3745– 3761 (2005).

17. Goyal, P. et al. Structural and mechanistic insights into the bacterial amyloid secretion channel CsgG. Nature 516, 250–253 (2014).

18. Derrington, I. M. et al. Nanopore DNA sequencing with MspA. Proceedings of the National Academy of Sciences 107, 16060–16065 (2010).

19. Chanakul, W. et al. Large and Stable Nanopores Formed by Complement Component 9 for Characterizing Single Folded Proteins. ACS Nano 19, 5240–5252 (2025).

20. Huang, G. et al. Electro-Osmotic Vortices Promote the Capture of Folded Proteins by PlyAB Nanopores. Nano Lett. 20, 3819–3827 (2020).

21. Huang, G. et al. PlyAB Nanopores Detect Single Amino Acid Differences in Folded Haemoglobin from Blood. Angewandte Chemie International Edition 61, e202206227 (2022).

22. Yusko, E. C. et al. Real-time shape approximation and fingerprinting of single proteins using a nanopore. Nat. Nanotechnol. 12, 360–367 (2017).

23. Chen, X. et al. Nanopore single-molecule analysis of biomarkers: Providing possible clues to disease diagnosis. TrAC Trends in Analytical Chemistry 162, 117060 (2023).

24. Yusko, E. C. et al. Controlling protein translocation through nanopores with bio-inspired fluid walls. Nat. Nanotechnol. 6, 253–260 (2011).

25. Houghtaling, J., List, J. & Mayer, M. Nanopore-Based, Rapid Characterization of Individual Amyloid Particles in Solution: Concepts, Challenges, and Prospects. Small 14, 1802412 (2018).

26. Yusko, E. C. et al. Single-Particle Characterization of Aβ Oligomers in Solution. ACS Nano 6, 5909–5919 (2012).

27. Houghtaling, J. et al. Estimation of Shape, Volume, and Dipole Moment of Individual Proteins Freely Transiting a Synthetic Nanopore. ACS Nano 13, 5231–5242 (2019).

28. Awasthi, S., Ying, C., Li, J. & Mayer, M. Simultaneous Determination of the Size and Shape of Single α-Synuclein Oligomers in Solution. ACS Nano 17, 12325–12335 (2023).

29. Prangkio, P., Yusko, E. C., Sept, D., Yang, J. & Mayer, M. Multivariate Analyses of Amyloid-Beta Oligomer Populations Indicate a Connection between Pore Formation and Cytotoxicity. PLoS One 7, e47261 (2012).

30. Ying, Y.-L. et al. Nanopore-based technologies beyond DNA sequencing. Nat. Nanotechnol. 17, 1136–1146 (2022).

31. Ying, Y.-L. & Long, Y.-T. Nanopore-Based Single-Biomolecule Interfaces: From Information to Knowledge. J. Am. Chem. Soc. 141, 15720–15729 (2019).

32. Jiang, J. et al. Protein nanopore reveals the renin–angiotensin system crosstalk with single-amino-acid resolution. Nat. Chem. 15, 578–586 (2023).

33. Galenkamp, N. S., Biesemans, A. & Maglia, G. Directional conformer exchange in dihydrofolate reductase revealed by single-molecule nanopore recordings. Nat. Chem. 12, 481–488 (2020).

34. Acharjee, M. C. et al. Tau and tubulin protein aggregation characterization by solid-state nanopore method and atomic force microscopy. J. Appl. Phys. 133, 024701 (2023).

35. Sandler, S. E. et al. Multiplexed Digital Characterization of Misfolded Protein Oligomers via Solid-State Nanopores. J. Am. Chem. Soc. 145, 25776–25788 (2023).

36. Wang, H.-Y., Ying, Y.-L., Li, Y., Kraatz, H.-B. & Long, Y.-T. Nanopore Analysis of β-Amyloid Peptide Aggregation Transition Induced by Small Molecules. Anal. Chem. 83, 1746–1752 (2011).

37. Meyer, N. et al. Solid-state and polymer nanopores for protein sensing: A review. Adv. Colloid Interface Sci. 298, 102561 (2021).

38. Tavassoly, O., Kakish, J., Nokhrin, S., Dmitriev, O. & Lee, J. S. The use of nanopore analysis for discovering drugs which bind to α-synuclein for treatment of Parkinson’s disease. Eur. J. Med. Chem. 88, 42–54 (2014).

39. Giamblanco, N. et al. Mechanisms of Heparin-Induced Tau Aggregation Revealed by a Single Nanopore. ACS Sens. 5, 1158–1167 (2020).

40. Yu, R.-J. et al. Single molecule sensing of amyloid-β aggregation by confined glass nanopores. Chem. Sci. 10, 10728–10732 (2019).

41. Niedzwiecki, D. J., Iyer, R., Borer, P. N. & Movileanu, L. Sampling a Biomarker of the Human Immunodeficiency Virus across a Synthetic Nanopore. ACS Nano 7, 3341–3350 (2013).

42. Luo, X. & Davis, J. J. Electrical biosensors and the label free detection of protein disease biomarkers. Chem. Soc. Rev. 42, 5944 (2013).

43. Rusling, J. F., Kumar, C. V., Gutkind, J. S. & Patel, V. Measurement of biomarker proteins for point-of-care early detection and monitoring of cancer. Analyst 135, 2496 (2010).

44. Krainer, G. et al. Direct digital sensing of protein biomarkers in solution. Nat. Commun. 14, 653 (2023).

45. Fragasso, A., Schmid, S. & Dekker, C. Comparing Current Noise in Biological and Solid-State Nanopores. ACS Nano 14, 1338–1349 (2020).

46. Straathof, S. et al. Protein Sizing with 15 nm Conical Biological Nanopore YaxAB. ACS Nano 17, 13685–13699 (2023).

47. Fukumoto, H. et al. High-molecular-weight β-amyloid oligomers are elevated in cerebrospinal fluid of Alzheimer patients. The FASEB Journal 24, 2716–2726 (2010).

48. Dujardin, S. et al. Tau molecular diversity contributes to clinical heterogeneity in Alzheimer’s disease. Nat. Med. 26, 1256–1263 (2020).

49. Cárdenas-Aguayo, M. del C., Gómez-Virgilio, L., DeRosa, S. & Meraz-Ríos, M. A. The Role of Tau Oligomers in the Onset of Alzheimer’s Disease Neuropathology. ACS Chem. Neurosci. 5, 1178–1191 (2014).

50. Mannini, B. et al. Toxicity of Protein Oligomers Is Rationalized by a Function Combining Size and Surface Hydrophobicity. ACS Chem. Biol. 9, 2309–2317 (2014).

51. van Pee, K. et al. CryoEM structures of membrane pore and prepore complex reveal cytolytic mechanism of Pneumolysin. Elife 6, e23644 (2017).

52. Peraro, M. D. & van der Goot, F. G. Pore-forming toxins: ancient, but never really out of fashion. Nat. Rev. Microbiol. 14, 77–92 (2016).

53. Ward, S. M., Himmelstein, D. S., Lancia, J. K. & Binder, L. I. Tau oligomers and tau toxicity in neurodegenerative disease. Biochem. Soc. Trans. 40, 667–671 (2012).

54. Meraz-Ríos, M. A., Lira-De León, K. I., Campos-Peña, V., De Anda-Hernández, M. A. & Mena- López, R. Tau oligomers and aggregation in Alzheimer’s disease. J. Neurochem. 112, 1353– 1367 (2010).

55. Lasagna-Reeves, C. A. et al. Alzheimer brain-derived tau oligomers propagate pathology from endogenous tau. Sci. Rep. 2, 700 (2012).

56. Mietelska-Porowska, A., Wasik, U., Goras, M., Filipek, A. & Niewiadomska, G. Tau Protein Modifications and Interactions: Their Role in Function and Dysfunction. Int. J. Mol. Sci. 15, 4671–4713 (2014).

57. Drubin, D. G. & Kirschner, M. W. Tau protein function in living cells. J. Cell Biol. 103, 2739– 2746 (1986).

58. Kovacs, G. G. Invited review: Neuropathology of tauopathies: principles and practice. Neuropathol. Appl. Neurobiol. 41, 3–23 (2015).

59. Limorenko, G. & Lashuel, H. A. Revisiting the grammar of Tau aggregation and pathology formation: how new insights from brain pathology are shaping how we study and target Tauopathies. Chem. Soc. Rev. 51, 513–565 (2022).

60. El Mammeri, N., Dregni, A. J., Duan, P., Wang, H. K. & Hong, M. Microtubule-binding core of the tau protein. Sci. Adv. 8, eabo4459 (2022).

61. Fitzpatrick, A. W. P. et al. Cryo-EM structures of tau filaments from Alzheimer’s disease. Nature 547, 185–190 (2017).

62. von Bergen, M., Barghorn, S., Biernat, J., Mandelkow, E.-M. & Mandelkow, E. Tau aggregation is driven by a transition from random coil to beta sheet structure. Biochimica et Biophysica Acta (BBA) - Molecular Basis of Disease 1739, 158–166 (2005).

63. Giamblanco, N. et al. Mechanisms of Heparin-Induced Tau Aggregation Revealed by a Single Nanopore. ACS Sens. 5, 1158–1167 (2020).

64. Monien, B. H., Apostolova, L. G. & Bitan, G. Early diagnostics and therapeutics for Alzheimer’s disease – how early can we get there? Expert Rev. Neurother. 6, 1293–1306 (2006).

65. Lo, C. H. et al. Targeting the ensemble of heterogeneous tau oligomers in cells: A novel small molecule screening platform for tauopathies. Alzheimer’s & Dementia 15, 1489–1502 (2019).

66. Mamun, A., Uddin, M., Mathew, B. & Ashraf, G. Toxic tau: structural origins of tau aggregation in Alzheimer’s disease. Neural Regen. Res. 15, 1417 (2020).

67. Puangmalai, N. et al. Internalization mechanisms of brain-derived tau oligomers from patients with Alzheimer’s disease, progressive supranuclear palsy and dementia with Lewy bodies. Cell Death Dis. 11, 314 (2020).

68. Berger, Z. et al. Accumulation of Pathological Tau Species and Memory Loss in a Conditional Model of Tauopathy. The Journal of Neuroscience 27, 3650–3662 (2007).

69. Sanders, D. W. et al. Distinct Tau Prion Strains Propagate in Cells and Mice and Define Different Tauopathies. Neuron 82, 1271–1288 (2014).

70. Vaquer-Alicea, J. & Diamond, M. I. Propagation of Protein Aggregation in Neurodegenerative Diseases. Annu. Rev. Biochem. 88, 785–810 (2019).

71. Baker, N. A., Sept, D., Joseph, S., Holst, M. J. & McCammon, J. A. Electrostatics of nanosystems: Application to microtubules and the ribosome. Proceedings of the National Academy of Sciences 98, 10037–10041 (2001).

72. Vögele, M. et al. Membrane perforation by the pore-forming toxin pneumolysin. Proceedings of the National Academy of Sciences 116, 13352–13357 (2019).

73. van Pee, K., Mulvihill, E., Müller, D. J. & Yildiz, Ö. Unraveling the Pore-Forming Steps of Pneumolysin from *Streptococcus pneumoniae*. Nano Lett. 16, 7915–7924 (2016).

74. Wolfmeier, H. et al. Active release of pneumolysin prepores and pores by mammalian cells undergoing a Streptococcus pneumoniae attack. Biochimica et Biophysica Acta (BBA) - General Subjects 1860, 2498–2509 (2016).

75. Tilley, S. J., Orlova, E. V., Gilbert, R. J. C., Andrew, P. W. & Saibil, H. R. Structural Basis of Pore Formation by the Bacterial Toxin Pneumolysin. Cell 121, 247–256 (2005).

76. Larpin, Y. et al. Bacterial pore-forming toxin pneumolysin: Cell membrane structure and microvesicle shedding capacity determines differential survival of immune cell types. The FASEB Journal 34, 1665–1678 (2020).

77. Shen, Q. et al. Functionalized DNA-Origami-Protein Nanopores Generate Large Transmembrane Channels with Programmable Size-Selectivity. J. Am. Chem. Soc. 145, 1292–1300 (2023).

78. Yusko, E. C. et al. Real-time shape approximation and fingerprinting of single proteins using a nanopore. Nat. Nanotechnol. 12, 360–367 (2017).

79. Golibersuch, D. C. Observation of Aspherical Particle Rotation in Poiseuille Flow via the Resistance Pulse Technique. Biophys. J. 13, 265–280 (1973).

80. DeBlois, R. W., Uzgiris, E. E., Cluxton, D. H. & Mazzone, H. M. Comparative measurements of size and polydispersity of several insect viruses. Anal. Biochem. 90, 273–288 (1978).

81. Li, Y., Ying, C. & Mayer, M. Continuous, Low Latency Estimation of the Size and Shape of Single Proteins from Real-Time Nanopore Data. Anal. Chem. 98, 224–234 (2026).

82. Popov, K. I., Makepeace, K. A. T., Petrotchenko, E. V., Dokholyan, N. V. & Borchers, C. H. Insight into the Structure of the “Unstructured” Tau Protein. Structure 27, 1710–1715.e4 (2019).

83. Zhang, W. et al. Heparin-induced tau filaments are polymorphic and differ from those in Alzheimer’s and Pick’s diseases. Elife 8, e43584 (2019).

84. Zhu, H.-L. et al. Quantitative Characterization of Heparin Binding to Tau Protein. Journal of Biological Chemistry 285, 3592–3599 (2010).

85. Erickson, H. P. Size and Shape of Protein Molecules at the Nanometer Level Determined by Sedimentation, Gel Filtration, and Electron Microscopy. Biol. Proced. Online 11, 32–51 (2009).

86. Paul, S. S., Lyons, A., Kirchner, R. & Woodside, M. T. Quantifying Oligomer Populations in Real Time during Protein Aggregation Using Single-Molecule Mass Photometry. ACS Nano 16, 16462–16470 (2022).

87. Patterson, K. R. et al. Characterization of Prefibrillar Tau Oligomers in Vitro and in Alzheimer Disease. Journal of Biological Chemistry 286, 23063–23076 (2011).

88. Makrides, V. et al. Microtubule-dependent Oligomerization of Tau. Journal of Biological Chemistry 278, 33298–33304 (2003).

89. Lasagna-Reeves, C. A. et al. Identification of oligomers at early stages of tau aggregation in Alzheimer’s disease. The FASEB Journal 26, 1946–1959 (2012).

90. Gerson, J. E., Mudher, A. & Kayed, R. Potential mechanisms and implications for the formation of tau oligomeric strains. Crit. Rev. Biochem. Mol. Biol. 51, 482–496 (2016).

91. Drücker, P. et al. Pneumolysin-damaged cells benefit from non-homogeneous toxin binding to cholesterol-rich membrane domains. Biochimica et Biophysica Acta (BBA) - Molecular and Cell Biology of Lipids 1863, 795–805 (2018).

92. Wang, J., Terrasse, R., Bafna, J. A., Benier, L. & Winterhalter, M. Electrophysiological Characterization of Transport Across Outer-Membrane Channels from Gram-Negative Bacteria in Presence of Lipopolysaccharides. Angewandte Chemie International Edition 59, 8517–8521 (2020).

93. Cruickshank, C. C., Minchin, R. F., Le Dain, A. C. & Martinac, B. Estimation of the pore size of the large-conductance mechanosensitive ion channel of Escherichia coli. Biophys. J. 73, 1925–1931 (1997).

94. Crespo, R., Koudstaal, W. & Apetri, A. In Vitro Assay for Studying the Aggregation of Tau Protein and Drug Screening. Journal of Visualized Experiments 141, e58570 (2018).

95. Schneider, C. A., Rasband, W. S. & Eliceiri, K. W. NIH Image to ImageJ: 25 years of image analysis. Nat. Methods 9, 671–675 (2012).

