## Supplementary Information for "Pneumolysin nanopores with a 20 nm diameter characterize the size and shape of individual Tau oligomers"

[3] Present Address: Department of Biotechnology, National Institute of Pharmaceutical Education and Research Raebareli (NIPER-R), Lucknow, UP, India




#### Table of Contents


Supplementary Note 1: Estimation of pore inner diameter and number of monomers 2

Supplementary Note 2: Effect of pore cross-sectional geometry on pore diameter 3

Supplementary Note 3: Data analysis algorithm 4

Supplementary Note 4: Approximation of protein shape with an ellipsoid of rotation 6

Supplementary Note 5: Approximation of protein molecular weight from its radius 7

Supplementary Note 6: TEM Image Analysis of Tau oligomers 7

Supplementary Note 7: Event Frequency Calculation 8

Supplementary Note 8: Event frequency fitting 9

Supplementary Note 9: Concentration calculation of Tau oligomers 10

Supplementary Figures 11

Supplementary Table 25

References 26

***Supplementary Note 1: Estimation of inner pore diameter and number of monomers***

We calculated the equivalent cylindrical PLY pore diameter based on its conductance by comparing the baseline current with the current after single-pore insertion following the procedure reported by Chanakul *et al.*[1] for poly(C9) nanopores and equation (1) presented by Cruickshank *et al.*:[2]

$$d_{\mathrm{p}} = \frac{\rho_{\mathrm{el}}\, G}{\pi} \times \left(\frac{\pi}{2} + \sqrt{\frac{\pi^2}{4} + \frac{4\pi l_{\mathrm{p}}}{\rho_{\mathrm{el}}\, G}}\right) \tag{1}$$

In this equation:

$d_{\mathrm{p}}\ (\mathrm{m})$ — Inner diameter of a cylindrical nanopore,

$\rho_{\mathrm{el}}\ (\Omega \cdot \mathrm{m})$ — Electrical resistivity of the electrolyte buffer, we used an experimentally measured value of 4.75 $\Omega \cdot \mathrm{m}$ for the recording buffer with 500 mM NaCl and 50 mM Tris-HCl.

$G\ (\Omega^{-1})$ — Single channel conductance, determined as the difference between the baseline current and the open pore current at the applied voltage,

$l_{\mathrm{p}}\ (\mathrm{m})$ — Effective pore length. We determined it to be 9.5 nm (see **Supplementary Figure S4**) and assumed it to be constant for all pore diameters since all PLY pores are made up of the same PLY monomers.

We then used the determined pore diameter to estimate the number of PLY monomers comprising the pore using a geometric model described by Fennouri *et al.*:[3]

$$d_p = s\sqrt{\frac{1}{\pi}\left(\frac{n}{tan\left(\frac{\pi}{n}\right)} - \frac{(n-2)\pi}{2}\right)} \tag{2}$$

In this equation:

$d_{\mathrm{p}}\ (\mathrm{m})$ — Inner diameter of the nanopore, obtained experimentally using Equation (1),

$n$ — Number of monomers in the PLY pore,

$s$ (m) — Diameter of a cylindrical rod representing a PLY monomer.

The model assumes that the PLY pore is a circular arrangement of cylindrical rods (PLY monomers) oriented perpendicular to the surface of a planar lipid bilayer. These rods are positioned at the corner points of a regular polygon, with the diameter of each rod equal to the distance between the neighboring corner points.

We have previously demonstrated[2] that a Taylor expansion of Equation (2) provides an accurate estimation of pore diameter: as a function of the number of monomers:

$$d_p \approx s(0.318n - 0.784) \tag{3}$$

To determine the diameter $s$ of a cylindrical PLY monomer, we used Equation (3) and information from a recent cryo-EM structure[4] where the inner diameter of PLY was found to be d = 22 nm, and the pore consists of n = 42 monomers. Consequently, the diameter, s, of a cylindrical rod representing a PLY monomer in a PLY nanopore is approximately 2.02 nm. We used this value to estimate the number of monomers from the measured open pore conductance.

***Supplementary Note 2: Effect of pore cross-sectional geometry on pore diameter***

To evaluate the influence of pore shape on the determination of PLY pore diameter from single-channel conductance measurements, we compared the standard cylindrical pore model with pores having non-circular cross-sections while maintaining the same conductance. The cylindrical pore model including Hall access resistance on both sides of the membrane is given by: [1,2]

$$R = \frac{1}{\sigma}\left(\frac{4L}{\pi D^2} + \frac{1}{D}\right) \quad (4)$$

In this equation:

$R\ (\Omega)$ — Total pore resistance,

$D\ (m)$ — Inner diameter of a cylindrical nanopore,

$\sigma\ (S\ m^{-1})$— Conductivity of the electrolyte,

$L$— Effective pore length.

For the conditions used throughout this study, we assumed an effective pore length of $L = 9.5\ nm$ (**Supplementary Figure S8**) and an experimentally measured electrolyte resistivity of $\rho = 4.75\ \Omega m$ for the recording buffer containing 500 mM NaCl and 50 mM Tris-HCl. The corresponding conductivity is $\sigma = \frac{1}{\rho} = 0.211\ S\ m^{-1}$. To investigate the effect of pore shape, we considered a pore with a conductance corresponding to a cylindrical diameter of $D = 20\ nm$ and compared this idealized pore with pores possessing elliptical cross-sections. For an ellipse with major and minor axes $D_{max}$ and $D_{min}$ respectively, the area-equivalent diameter is,

$$D_A = \sqrt{D_{\max} D_{\min}} \quad (5)$$

which is the diameter of a circular pore having the same cross-sectional area. For a pore with a 2:1 elliptical cross-section ($Dmax = 2Dmin$), the access resistance differs slightly from that of a circular pore of identical area. Equating the total resistance of the elliptical pore to that of a cylindrical pore with $D = 20\ nm$ and $L = 9.5\ nm$ yields an area-equivalent diameter,

$$D_A = 19.74\ nm$$

The corresponding ellipse dimensions are,

$$D_{min} = 13.96\ nm$$

and $$D_{max} = 27.91\,nm$$

Therefore, the conductance-derived cylindrical diameter overestimates the area-equivalent diameter by

$$\frac{20.00 - 19.74}{19.74} \times 100\% \approx 1.3\%$$

Because both the pore resistance and the access resistance scale inversely with electrolyte conductivity, the shape-dependent correction is independent of electrolyte concentration provided that the conductivity is spatially uniform throughout the pore and access regions. Consequently, the result remains unchanged for measurements performed in either 0.5 M or 1 M NaCl. For comparison, the same analysis was performed for a more elongated 3:1 ellipse ($D_{\max} = 3D_{\min}$). In this case, the area-equivalent diameter is approximately 19.37 nm, corresponding to a diameter overestimation of approximately 3.3%.

These calculations demonstrate that the conductance-derived pore diameter is only weakly affected by moderate deviations from circularity. Thus, for the typical dimensions of PLY pores with a diameter close to 20 nm and a length of 9.5 nm, the diameter obtained from the experimentally determined single-channel conductance using a cylindrical pore model is less than 1.5% larger than the area-equivalent diameter of a pore with a 2:1 elliptical cross-section. Even for a substantially elongated 3:1 elliptical pore, the discrepancy remains small at approximately 3.3%. Thus, under the conditions considered here, conductance measurements primarily reflect the effective cross-sectional area of the pore and are only weakly sensitive to deviations from circular geometry, such that a cylindrical pore model results in excellent estimates of pore diameter for all but the most extreme non-circular cross-sections. Cryo-EM studies of PLY pores in lipid membranes revealed that the majority of PLY pores assembled from a closed ring of PLY monomers have cross-sections that are either closed to perfect circles or deviate less from a perfect circle than a 2:1 elliptical cross-section.[5]

***Supplementary Note 3: Data analysis algorithm***

We developed data analysis software to obtain excluded volume $\Lambda$ and length-to-diameter ratio $m$ of a target analyte from nanopore recordings.[1]

The analysis consists of three sequential steps:

1. Baseline search. In the first step, the recording $x$, composed of $W$ samples are imported and digitally filtered using a Gaussian low-pass filter with a desired cutoff frequency. The data are subsequently processed with a custom-made baseline search algorithm that operates as follows:

1) The filtered recording $x^{\mathrm{f}}$, with size $W$, is subdivided into $N+1$ segments, $N$ containing $M$ samples each and one containing the remaining $W-NM$ samples. The value of $M$ is set by the user and should always be M >100 and at least 30 times longer than the longest translocation event.
2) An empty array $x^{\mathrm{b}}$ with length $W$ is initialized, which will be filled with the estimated baseline values. For clarity, the $j^{\mathrm{th}}$ element of an array $x$ is indicated as $x[j]$.
3) The mean ($\bar{y}_{\mathrm{i}}$) and the standard deviation ($\sigma_{\mathrm{i}}$) of each segment $y_i$ is calculated.
4) The minimum standard deviation $\sigma_{\mathrm{min}}$is identified as the minimum value among the standard deviations of the $M$-sized segments $y_i$.
5) A threshold parameter $\tau > 1$ is defined (typically 1.2< $\tau$ <1.6).
6) For each segment $y_i$, with $i = 1 \ldots N$, the standard deviation $\sigma_{\mathrm{i}}$ is compared to $\tau\sigma_{\mathrm{min}}$ with two possible outcomes:
   A) if $\sigma_{\mathrm{i}} \leq \tau\sigma_{\mathrm{min}}$, the elements of $x^{\mathrm{b}}$ comprised between $x^{\mathrm{b}}[(i-1)M]$ and $x^{\mathrm{b}}[iM]$ are filled with the average value $\bar{y}_{\mathrm{i}}$
   B) if $\sigma_{\mathrm{i}} > \tau\sigma_{\mathrm{min}}$, $y_i$ is subdivided into 10 samples, $z_k^i$ ($k = 1 \ldots 10$) of length M/10. The elements of $x^{\mathrm{b}}$ comprised between $x^{\mathrm{b}}[(i-1)M + M(k-1)/10]$ and $x^{\mathrm{b}}[(i-1)M + Mk/10]$ are filled with the average value of the $z_k^i$ segment, $\bar{z}_k^i$.
7) The baseline is finally completed by filling the last $W-NM$ elements of $x^{\mathrm{b}}$ with the mean value of the last $W-NM$ elements of $x^{\mathrm{f}}$.

The baseline trace $x^{\mathrm{b}}$ is important because it allows tracking step changes in the baseline current and, hence, changes in the diameter of the nanopore over time.

2. Event detection. After determining the baseline trace, the normalized ($\Delta I/I_0$) trace, $x^{\mathrm{n}}$ is calculated as $x^{\mathrm{n}} = (x^{\mathrm{f}} - x^{\mathrm{b}})/x^{\mathrm{b}}$ and processed with the following threshold-based event detection algorithm:

1) The trace $x^{\mathrm{n}}$ is subdivided into $F+1$ segments, $F$ containing $G$ samples each and one containing the remaining $W-FG$ samples.
2) For each $G$-sized segment $q_i$, with $i = 1 \ldots F$, the mean $\bar{q}_i$ and the standard deviation $\sigma_i$ are computed. Two thresholds are calculated, $T_{\mathrm{s}} = \bar{q} + u\sigma_i$, where $u > 1$ (typically $u > 4$), and $T_{\mathrm{e}} = \bar{q} + \sigma_i$.
3) A preliminary search is performed to identify at which position in $q_i$ the signal first exceeds $T_{\mathrm{s}}$ ($e_{\mathrm{s}}$, event start) and then falls below $T_{\mathrm{e}}$ ($e_e$, event end). For each event, the end and start position of the events in the trace $x^{\mathrm{n}}$ are calculated to include part of the baseline as $E_{\mathrm{s}} = e_{\mathrm{s}} + (i-1)G - 40$ and $E_{\mathrm{e}} = e_{\mathrm{e}} + (i-1)G + 40$, and stored (extended events).

4) The detected events are subsequently refined, as a simple threshold-based search could fragment a single translocation into multiple short events, or interpret noise spikes as translocation events:
   A) All samples in $x^{\mathrm{n}}$ with value smaller than $T_{\mathrm{s}}$ are stored in an array $w$ of size $R$.
   B) A matrix $B_{(R-Q)\times Q}$ is constructed by stacking Q delayed copies of $w$ , such that the j[th] column of *B* contains the elements of $w$ comprised between $w[j]$ and $w[R-Q+j]$.
   C) The standard deviation of each row of *B* is calculated, and the minimum value obtained is defined $\sigma_0$.
   D) Each extended event is fitted using two Gaussian peaks having the same standard deviation $\sigma_0$ and centered at positions $L_0$ and $L_1 (L_1 > L_0)$. $L_0$ and $L_1$ represent an estimate of the baseline and of the event amplitude, respectively.
   E) The event is fitted with a 2-level step-fitting algorithm using $L_0$ and $L_1$ as levels.
   F) The portion of the event which is best fitted by $L_1$ is considered as the refined event. If the duration of the event exceeds a user-defined minimum duration, the event is accepted and stored.

3. Event analysis. The refined resistive pulses are analyzed to determine the minimum and the maximum intensity values $(dI_{\min}, dI_{\max})$, which are needed to determine the volume $(\Lambda)$ and the ellipsoid-equivalent shape $(m)$ of the analyte particles. Since the measurements are affected by noise, the absolute maximum and minimum of the event trace are not good estimates of $dI_{\min}$ and $dI_{\max}$. Instead, we developed the following method:

1) A matrix $B_{(R-Q)\times Q}$ is constructed as described above. The 20[th] and the 80[th] percentile of each row of $B$ are calculated and stored in two arrays, $P_{20}$ and $P_{80}$, with sizes $(R-Q)$.
2) The minimum value of the array obtained from the elementwise difference $P_{80}-P_{20}$ is labelled $dI_0$.
3) The 20[th] and the 80[th] percentile of each event, $dI_{20}$ and $dI_{80}$ is computed.
4) $dI_{\min}$ and $dI_{\max}$ are calculated as $dI_{\min} = dI_{20} + dI_0/2$ and $dI_{\max} = dI_{80} - dI_0/2$.
5) $dI_{\min}$ and $dI_{\max}$ are used to compute $\Lambda$ and $m$ as described in by Yusko *et al.*[6] and Houghtaling *et al.*[7]

***Supplementary Note 4: Approximation of protein shape with an ellipsoid of revolution***

We developed software to derive reference values for excluded volume $\Lambda_{\mathrm{r}}$ and length-to-diameter ratio $m_{\mathrm{r}}$ of a target analyte from atomic coordinates (*.PDB file):[1]

1) Atomic coordinates of a target protein were extracted from a *.PDB file

2) A grid with lattice dimension *l* was constructed to fully incorporate the protein.
3) The number of cells $N_C$ in the grid, at least one atom of the protein was counted.
4) The volume of the protein was determined as $\Lambda_r = N_C l^3$.
5) All coordinates were rescaled by subtracting their average values ($x_i' = x_i - \bar{x}$, $y_i' = y_i - \bar{y}, z_i' = z_i - \bar{z}$).
6) The equation of an ellipsoid of rotation with volume $\Lambda_r$ was calculated as a function of the axis ratio $m_r$, the coordinates of the center ($x_0, y_0$ and $z_0$), and the three Euler angles around the ellipsoid axis ($\alpha, \beta$ and $\gamma$).
7) $x_0$, $y_0$, $z_0$, $\alpha$, $\beta$, $\gamma$, and $m_r$ were optimized using a non-linear fitting procedure based on a search method to maximize the number of atoms of the protein enclosed within the ellipsoid.

***Supplementary Note 5: Approximation of protein molecular weight from its radius***

We calculated the molecular weight of a single protein from its radius, assuming a perfectly spherical shape of the particles, using Equations 3 and 4.[8]

$$r \approx 0.066 * (M.W.)^{1/3} \quad (3)$$

$$V = \frac{4}{3} * \pi * r^3 \quad (4)$$

$$V(nm^3) \approx 0.0012 \left(\frac{nm^3}{Da}\right) MW(Da) \quad (5)$$

Where,

$r$ is the radius of the protein, assuming a spherical model for its shape, nm.

$M.W.$ is the molecular weight of protein, Da.

$V$ is the volume-based molecular weight, nm$^3$.

***Supplementary Note 6: TEM image analysis of Tau oligomers***

We analyzed the transmission electron microscopy (TEM) micrographs of Tau oligomers during different stages of aggregation using the ImageJ (Fiji) software package (see **Supplementary Figure S14**). The analysis consisted of the following steps:

1) *Image preprocessing*: Raw TEM micrographs were imported, and image contrast was optimized through manual brightness and contrast adjustment. A mean filter (radius 2 pixels) was applied to reduce background noise and enhance particle visibility.
2) *Scale calibration*: The image scale was calibrated using the microscope-provided scale bar by setting the corresponding pixel-to-nanometer ratio within ImageJ.

3) *Particle measurement*: Individual Tau oligomers were manually selected using the oval selection tool in ImageJ. The projected area of each particle was measured, and the apparent diameter was calculated as the area-equivalent circular diameter. Only well-resolved, non-overlapping particles were included in the analysis. The compiled measurements from three independent micrographs (*N* = 3) were used to construct the size distribution of Tau oligomers.
4) *Estimation of protein molecular weight*: Flattening of proteins and protein assemblies during negative staining and air-drying for transmission electron microscopy (TEM) is a well-documented phenomenon.[9–13] The dehydration and adsorption of particles onto the TEM grid frequently lead to substantial axial compression, resulting in reduced apparent heights and distorted morphologies. Multiple studies have shown that negatively stained protein oligomers, amyloid assemblies, and other soft biomolecular complexes often exhibit significant flattening relative to their hydrated dimensions, with reported reductions in height ranging from ~30% to ~70% depending on particle size, rigidity, and staining conditions.[9–13] These observations indicate that lateral dimensions measured from negatively stained TEM images should be interpreted with caution, particularly when inferring volumetric or mass-related properties of flexible protein assemblies.

   Guided by our independent measurements from nanopore resistive-pulse sensing and mass photometry, we applied a 40% reduction in the measured TEM diameter of the particles to approximate their hydrated, native dimensions. Further, we estimated the molecular weight distribution of Tau oligomers from their radii, assuming a spherical particle shape as described in **Supplementary Note 5**.

***Supplementary Note 7: Calculation of Event Frequency***

We analyzed the event frequency of Tau oligomers from the nanopore recordings using a peak-area–based quantification workflow. Each ionic current trace was fitted with a multi-component Gaussian model to obtain the area $A_i$ corresponding to each oligomeric population $i$. The total number of events $N_i$ attributed to population $i$ was calculated according to

$$N_i = A_i \times N_{total} \tag{6}$$

where $N_{total}$ denotes the total number of events detected in the recording. The event frequency $f_i$ for population $i$ was then determined as

$$f_i = \frac{N_i}{t_{rec}} \tag{7}$$

where $t_{rec}$ is the total recording duration. This procedure ensured consistent quantification of all oligomeric populations across different aggregation time points.

***Supplementary Note 8: Fitting of Time-Dependent Aggregation of Tau with a Kinetic Model***

Event frequency data at different incubation times for different Tau oligomers were fitted using a Smoluchowski-like coagulation equation, where aggregation is treated as a set of irreversible binary collisions between all possible species present in solution:

$$\frac{d[O_i]}{dt} = -\sum_{j=1}^{\Omega} K_{ij}[O_i][O_j] + \frac{1}{2}\sum_{l=1}^{i-1} K_{l,l-i}[O_l][O_{l-i}] \tag{8}$$

In equation 8, $[O_i]$ is the molar concentration of oligomers with aggregation number (number of tau monomers) $i$, $K_{ij}$ and $K_{l,l-i}$ are a set of kinetic constants describing the consumption and creation of oligomers with an aggregation number $i$, respectively, while $\Omega$ is a cutoff parameter which sets the maximum oligomer size captured by the model (here we set $\Omega = 200$ since larger values increase the computation time without a real benefit on the kinetic description of the analyzed oligomers). Equation 8 is, in reality, a system of equations to be solved simultaneously for different $i$ values.

According to the collision theory,[14] the rate constants of each binary collision is expressed as

$$K_{ij} = 4\pi(D_i + D_j)(R_i + R_j)\frac{1}{1 - e^{-\frac{E_a}{k_B T}}}, \tag{9}$$

where $D_{i,j}$ $[\mathrm{dm}^2 \cdot \mathrm{s}^{-1}]$ are the diffusion coefficients of oligomers $i$ and $j$, $R_{i,j}$ $[\mathrm{dm}]$ are their respective radii, $T$ is temperature, $k_B$ is Boltzmann's constant and $E_a$ is an energy barrier, which is the only fitting parameter of the model and is assumed to be the same irrespectively of the size of the oligomers.

The radii of the oligomers are calculated from the radius of the monomer ($R_1 = 2.4$ nm) as:

$$R_i = i^{1/3} R_1 \tag{10}$$

While the diffusion coefficients are calculated from Stoke-Einstein's relation:

$$D_i = \frac{k_B T}{6\pi R_i \eta} \tag{11}$$

Where $\eta = 10^{-3}$ Pa·s approximates the viscosity of the solvent.

To find the value of $E_a$ which best reproduces the aggregation kinetics of Tau oligomers, we employed the following minimization algorithm:

1. We initialized the algorithm with concentrations of dimers and monomers matching those found experimentally at t=0
2. We integrated the system of equations 8 for i=1…5 using the solve_ivp module of Python's Scipy, where we used a Runge-Kutta integration scheme of order four ('RK45') for different values. At this stage, we assumed $E_a = 0$, which represents a purely diffusion-driven aggregation.
3. To estimate how well the model describes the experimental data, we calculated a modified residual sum of squares as follow
$$\chi_m^2 = \sum_t \sum_i [y(i,t) - f(i,t)]^2 i \tag{12}$$
Which represents the sum of the squared differences between data ($y$) and model (f) for the different oligomer sizes $i$ at different time points $t$. The squared differences are multiplied by $i$ to give all oligomeric species the same weight in the fitting.
4. Finally, we integrated the system of equations for increasing values of $E_a$ to find the value that minimizes $\chi_m^2$.

For fitting the TEM data, we adopted the same procedure, but instead of comparing event frequencies, we compared the relative abundances of species of various sizes present in the sample. This was done at each time step by dividing the number of particles, either measured or calculated, by the sum of the numbers of particles from the monomer to the 15-mer. This operation was necessary to compare data obtained at different times, where the natural variability in the sample preparation procedure and small differences in the size of the imaged area could introduce a significant bias.

The fitting procedure yielded an activation barrier $E_a = 10.5\ k_B T$ for the nanopore data and $E_a = 10.7\ k_B T$ for the TEM data. These values are in excellent agreement and physically reasonable, as they indicate slow aggregation kinetics in line with the experimental observations. The fitting curves are plotted in **Figure 3** in the main text and in **Supplementary Figure S17**. While this model is oversimplified, it provides a reasonable fit to the data in **Figure 3f** and provides visual guidance on oligomer abundance from dimer to pentamer as a function of time.

***Supplementary Note 8: Determination of the Concentration of Tau Oligomers***

We determined the concentration of Tau oligomers by converting the event frequency distribution into absolute concentrations using the known total Tau concentration in monomer equivalents (7.22 μM). First, the relative frequency of each oligomeric population was obtained by normalizing its event frequency to the total event frequency of the recording. These relative abundances were then scaled by the total monomer-equivalent concentration to obtain the monomer contribution of each species. Finally, the monomer-equivalent values were converted into absolute oligomer concentrations by accounting for the stoichiometry of each species (*i.e.*, the number of monomers per oligomer).

***Supplementary Figures:***

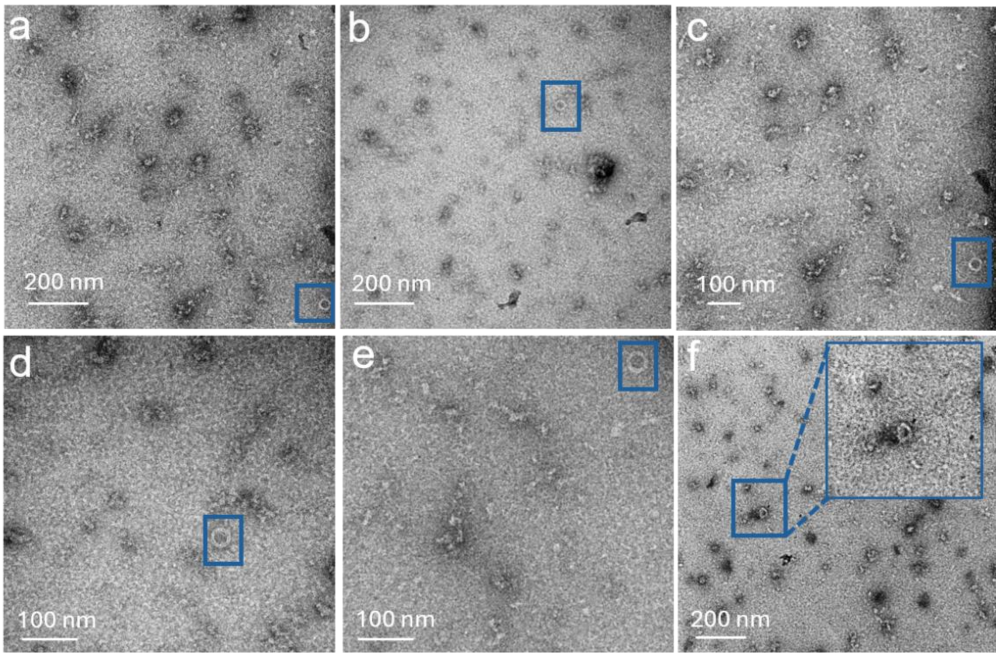


**Supplementary Figure S1. Transmission electron micrographs (TEMs) of PLY-amphipol incubated solution,** (**a-f**) representative independent experiments. The blue boxes indicate rings of the PLY-amphipol complex.

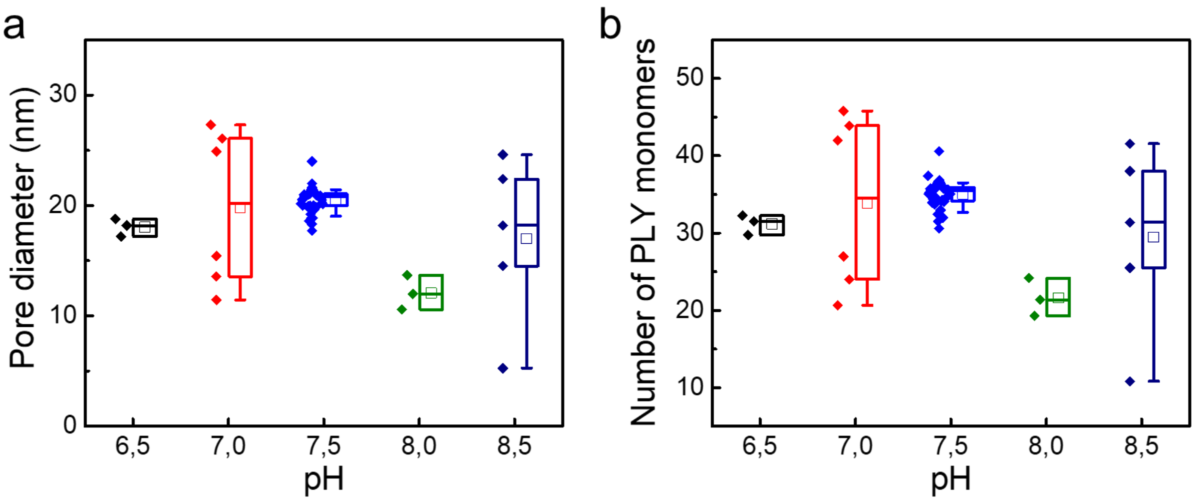


**Supplementary Figure S2. Estimation of the effective inner diameter of the PLY pores as a function of the pH value of the buffer solution.** (**a**) Inner diameter of PLY nanopores and (**b**) corresponding number of PLY monomers as a function of pH. For this calculation, we employed an effective pore length of 9.5 nm. The mean values are shown by the solid squares.

The box range corresponds to the 25th to 75th percentile, the whisker range corresponds to the 10th to 90th percentile.

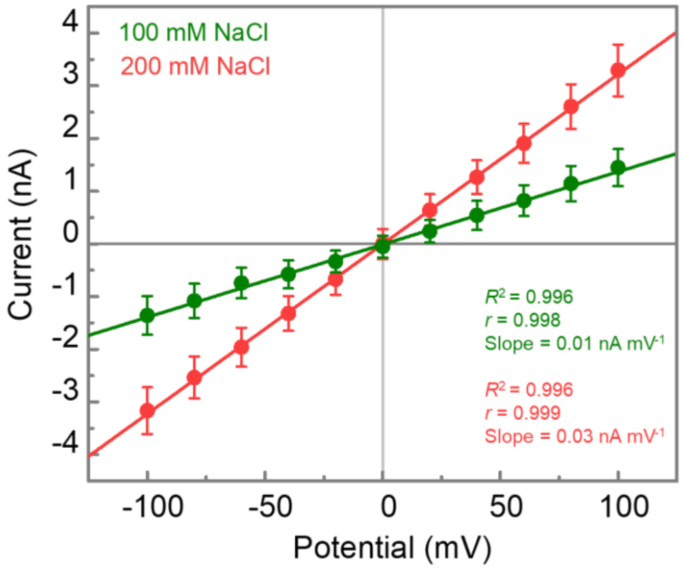


**Supplementary Figure S3. Current-voltage (I–V) curves of PLY pores in recording buffer with relatively low ionic strength.** The recording buffer contained 50 mM Tris-HCl, pH 7.5 with 100 mM and 200 mM NaCl. Error bars represent the standard deviations calculated from a minimum of three repeats.

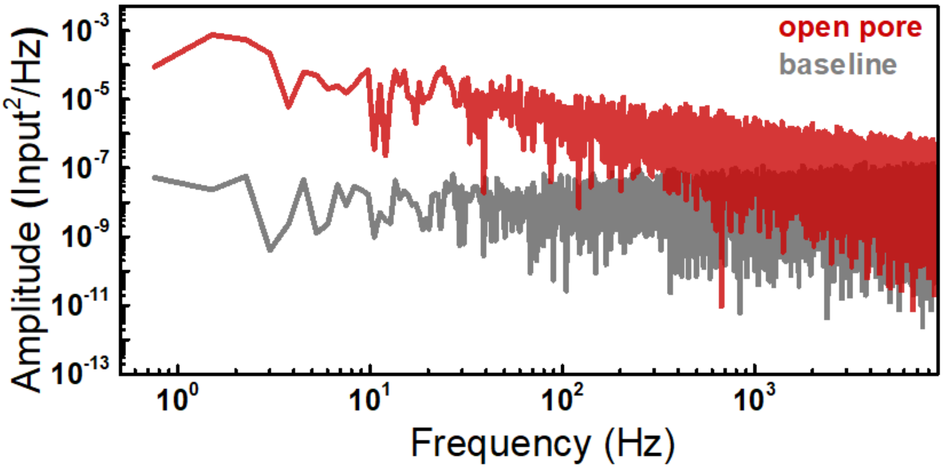


**Supplementary Figure S4**. **Noise analysis of PLY nanopores**. Comparison of power spectral densities (PSDs) from current before (baseline, grey) and after insertion of a single PLY pore (open pore, red) measured at +100 mV applied voltage in recording buffer containing 500 mM NaCl, 50 mM Tris-HCl, at pH 7.5. The PSD was determined from the current recordings that were collected with the maximum available sampling rate of 200 kHz of the amplifier without filtering.

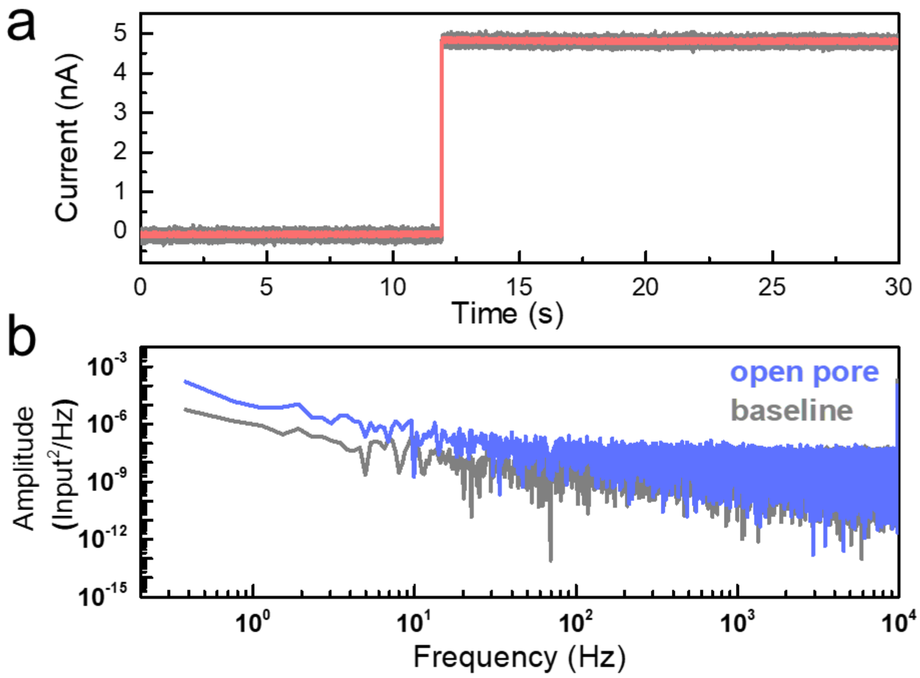


**Supplementary Figure S5. Characterization of PLY nanopores in the absence of cholesterol in the lipid bilayer. a)** Single-pore insertions of PLY into lipid bilayers that contained only DiphyPC lipids in the octane solution; no cholesterol was present. The current recordings were collected with a 200 kHz sampling rate (grey) and were filtered with a digital Gaussian low-pass filter with a cutoff frequency of 10 kHz (red). **b)** Comparison of power spectral densities (PSD) from current before (baseline, grey) and after insertion of a single PLY pore (open pore, blue). The PSD was determined from current recordings collected at a 200 kHz sampling rate, without filtering.

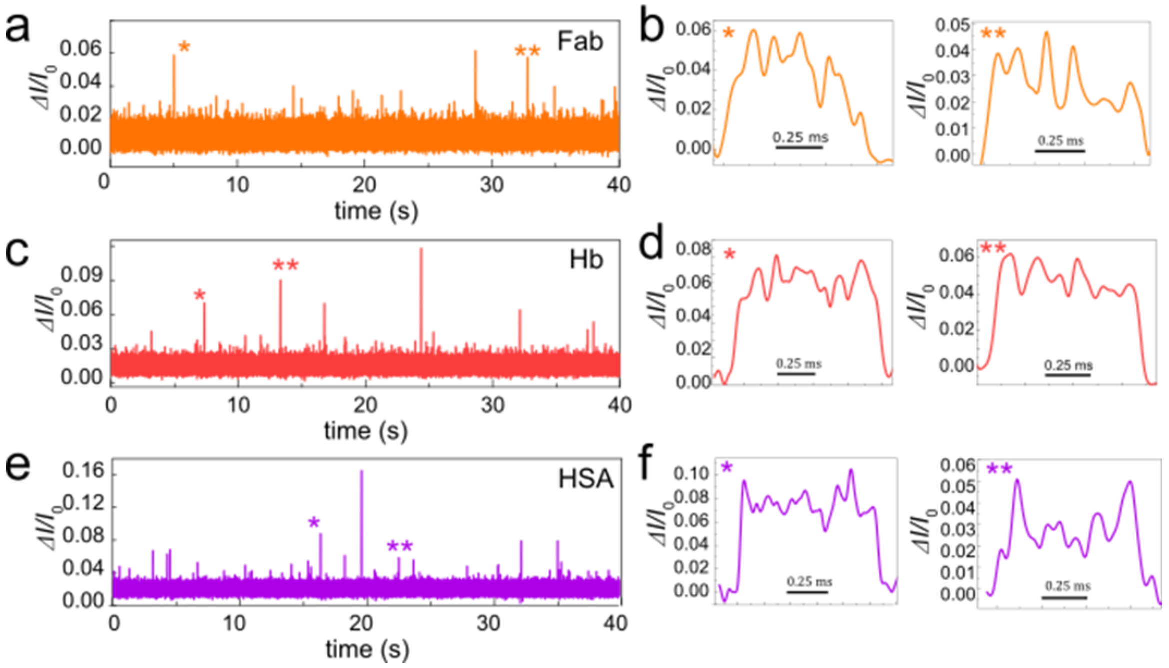


**Supplementary Figure S6. Baseline–corrected current recordings showing resistive pulses (upward spikes) and representative individual resistive pulses** measured with PLY nanopores in the presence of the following proteins: (**a, b**) Fab, (**c, d**) Hb, (**e, f**) HSA. The current recordings were collected in a buffer containing 500 mM NaCl, 0.2 μM amphipol, 50 mM Tris-HCl, pH 7.5, with a 200 kHz sampling rate and were filtered with a 10 kHz digital Gaussian low–pass filter for clarity of display.

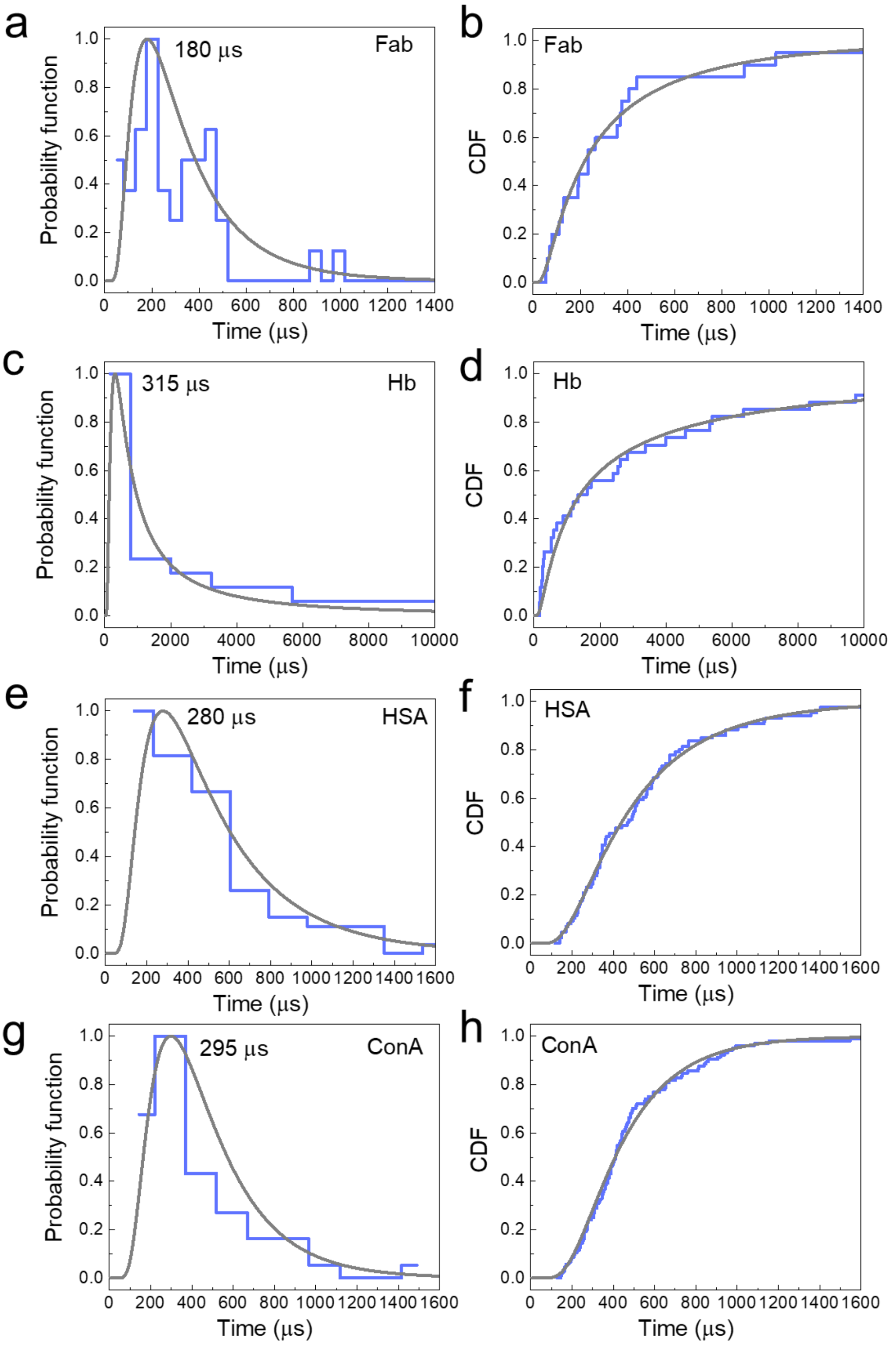


**Supplementary Figure S7. Distribution (PDF) of dwell times $t_d$ and the corresponding cumulative distribution function (CDF) of experimentally measured dwell times of resistive pulses from four proteins**. (**a**, **b**) Fab, (**c**, **d**) Hb, (**e**, **f**) HSA, (**g**, **h**) ConA. The distributions include the $t_d$ values of all detected resistive pulses that were longer than 20 µs and show that the most probable $t_d$ values for Fab, Hb, HSA, and ConA were 180 µs, 315 µs, 280 µs, and 295 µs, respectively. The CDFs were fitted to the experimental data and subsequently differentiated to obtain the corresponding grey curves of the probability density functions (PDFs) in a, c, e, and g; this procedure removes possible artifacts by the arbitrary choice of bin widths of the histograms.

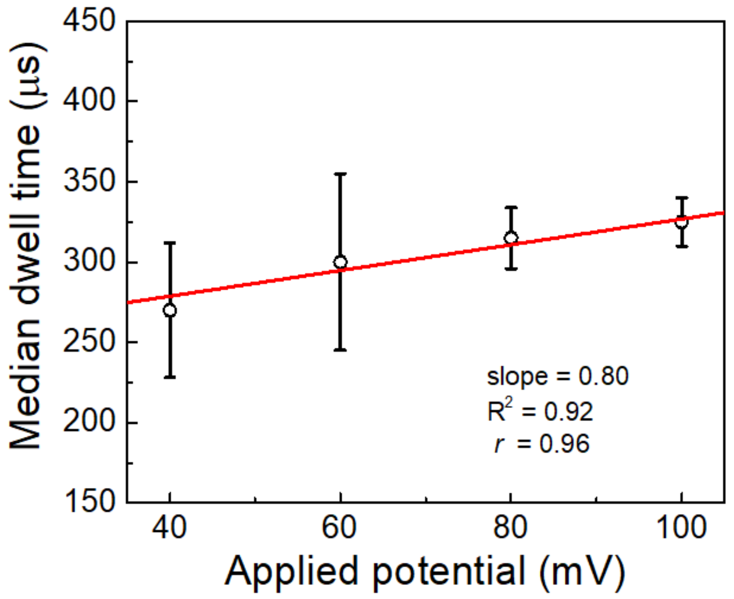


**Supplementary Figure S8. Voltage dependence of HSA dwell times through PLY nanopores.** Median dwell times of human serum albumin (HSA) resistive pulses measured at different applied potentials (−40 mV, −60 mV, −80 mV, and −100 mV). The analysis includes *N* = 33, 49, 248, and 481 events, respectively. The red line represents a linear least-squares regression fit ($R^2$ = 0.80) to the median dwell times. Error bars indicate the 5–95% confidence intervals. Experiments were performed in a buffer containing 500 mM NaCl 0.2 µM amphipol, 50 mM Tris-HCl, and pH 7.5.

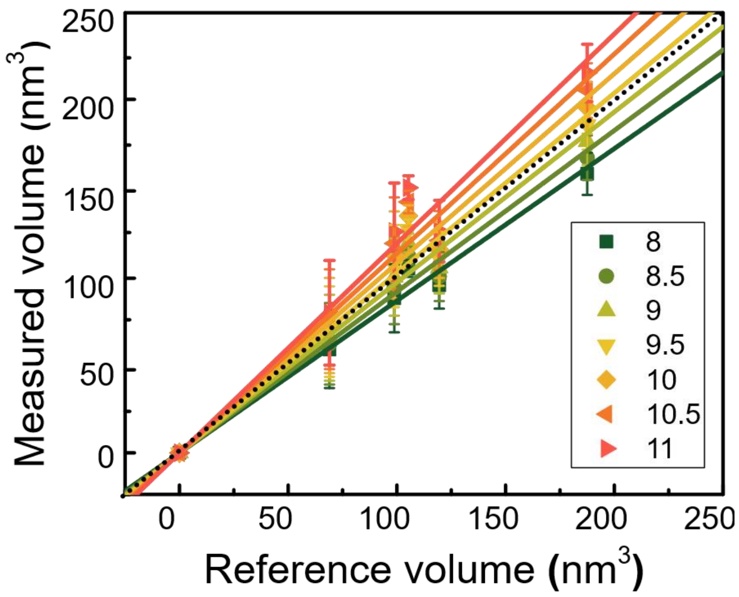


**Supplementary Figure S9. Experimental determination of the effective pore length of PLY pores by assuming various effective pore lengths for the determination of known protein volumes.** Comparison of excluded volume measured in PLY nanopore experiments with the reference volume determined from the atomic coordinates (*.PDB files). We fitted linear regressions with a zero intercept to ensure accurate analysis. The figure illustrates that an effective pore length of 9.5 nm most closely matches the ideal slope of 1.0 (black dotted line) for the measured values as a function of the reference values for five different proteins.

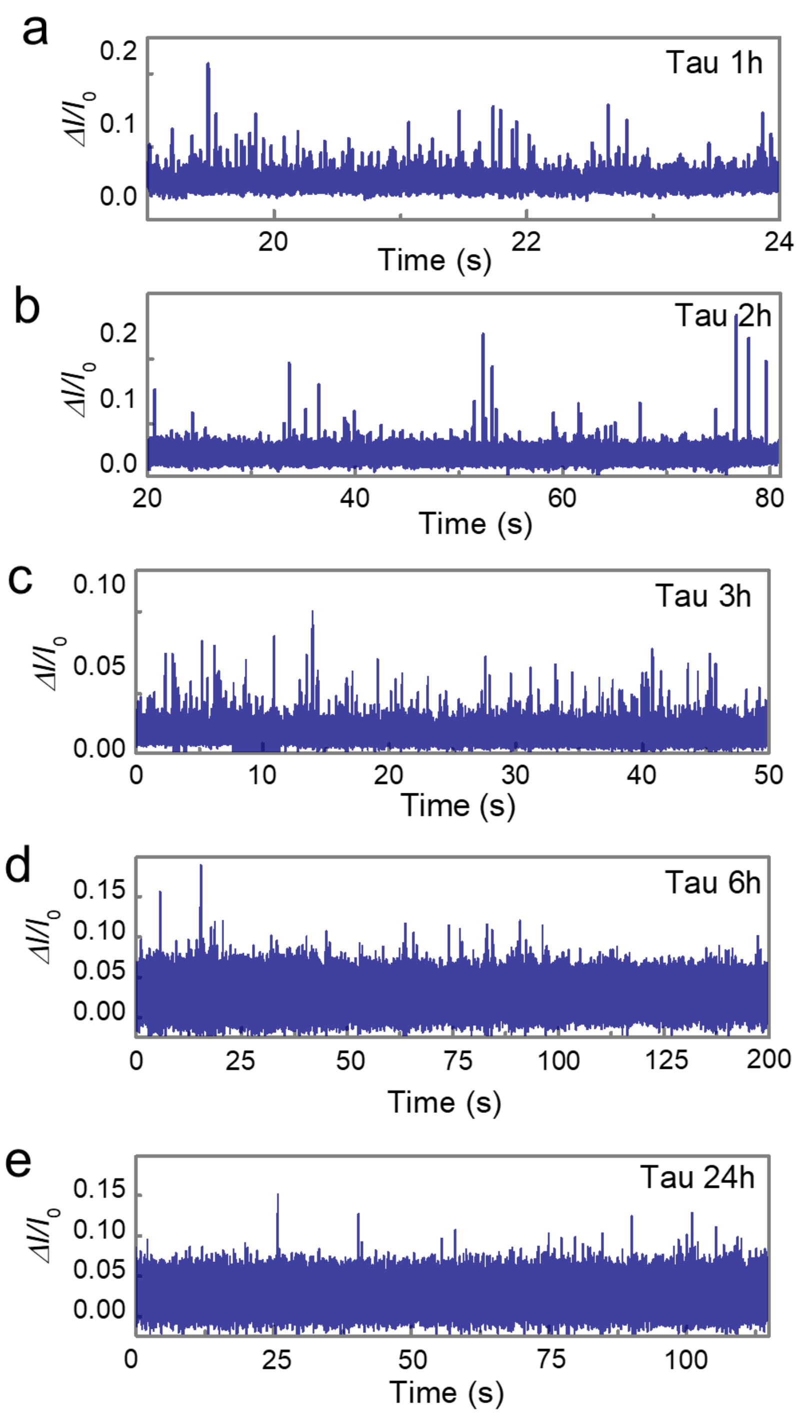


**Supplementary Figure S10. Baseline–corrected current recordings, showing resistive pulses in the presence of Tau oligomers (upward spikes)** (**a-e**) 1 h to 24 h after starting the aggregation**.** We used at least three independent PLY pores to record the size distribution of Tau oligomers at each time point. The current recordings were collected in a buffer containing 500 mM NaCl, 0.2 μM amphipol, 50 mM Tris-HCl, pH 7.5, with a 200 kHz sampling rate and were filtered with a 10 kHz digital Gaussian low–pass filter for clarity of display.

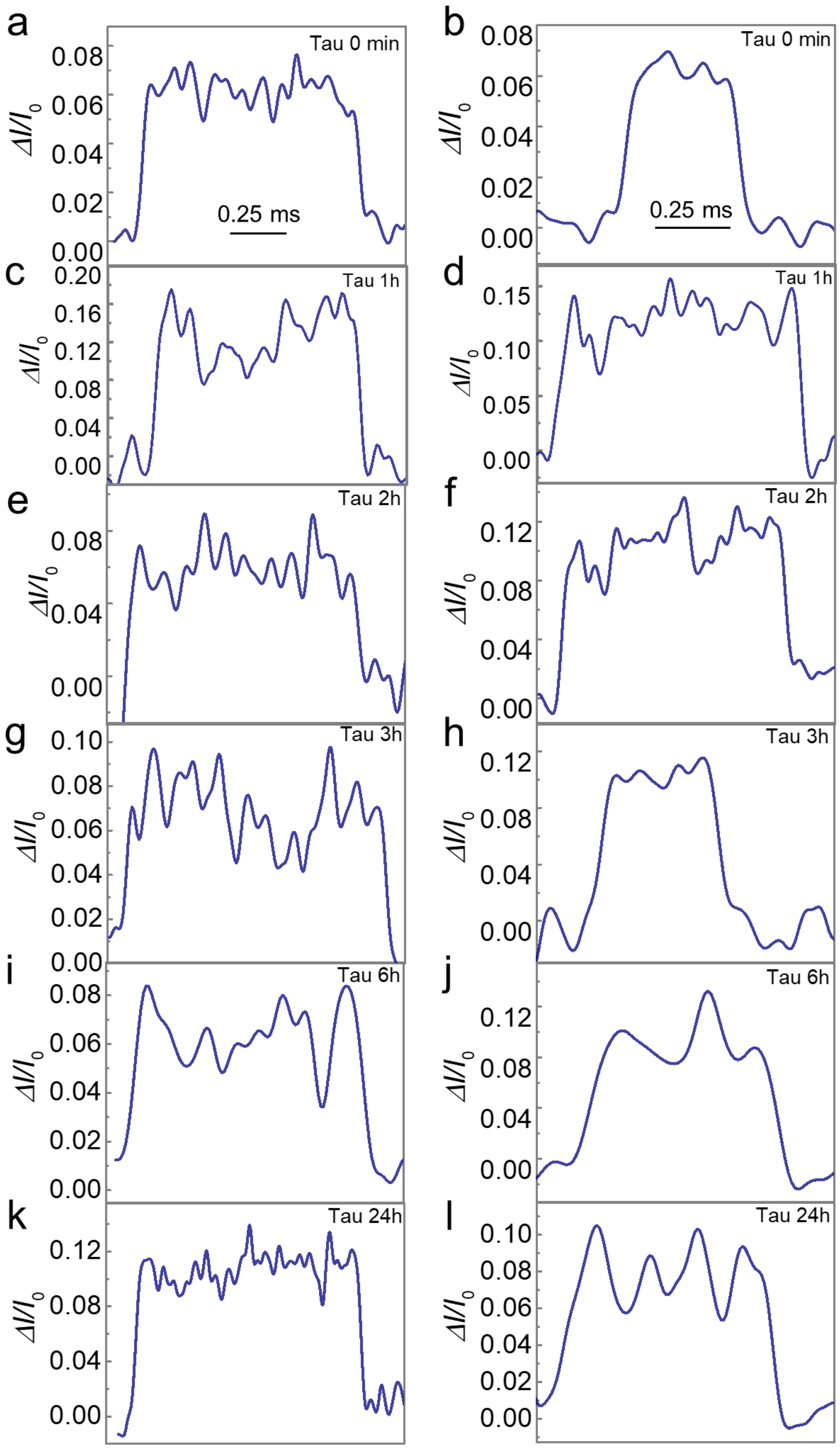

**Supplementary Figure S11. Representative individual resistive pulses in the presence of Tau oligomers** (**a, b**) 0 min, (**c, d**) 1 h, (**e, f**) 2 h, (**g, h**) 3 h, (**i, h**) 6 h, (**k, l**) 24 h time points during the aggregation**.** The current recordings were collected in a buffer containing 500 mM NaCl, 0.2 μM amphipol, 50 mM Tris-HCl, pH 7.5, with a 200 kHz sampling rate and were filtered with a 10 kHz digital Gaussian low–pass filter for clarity of display.

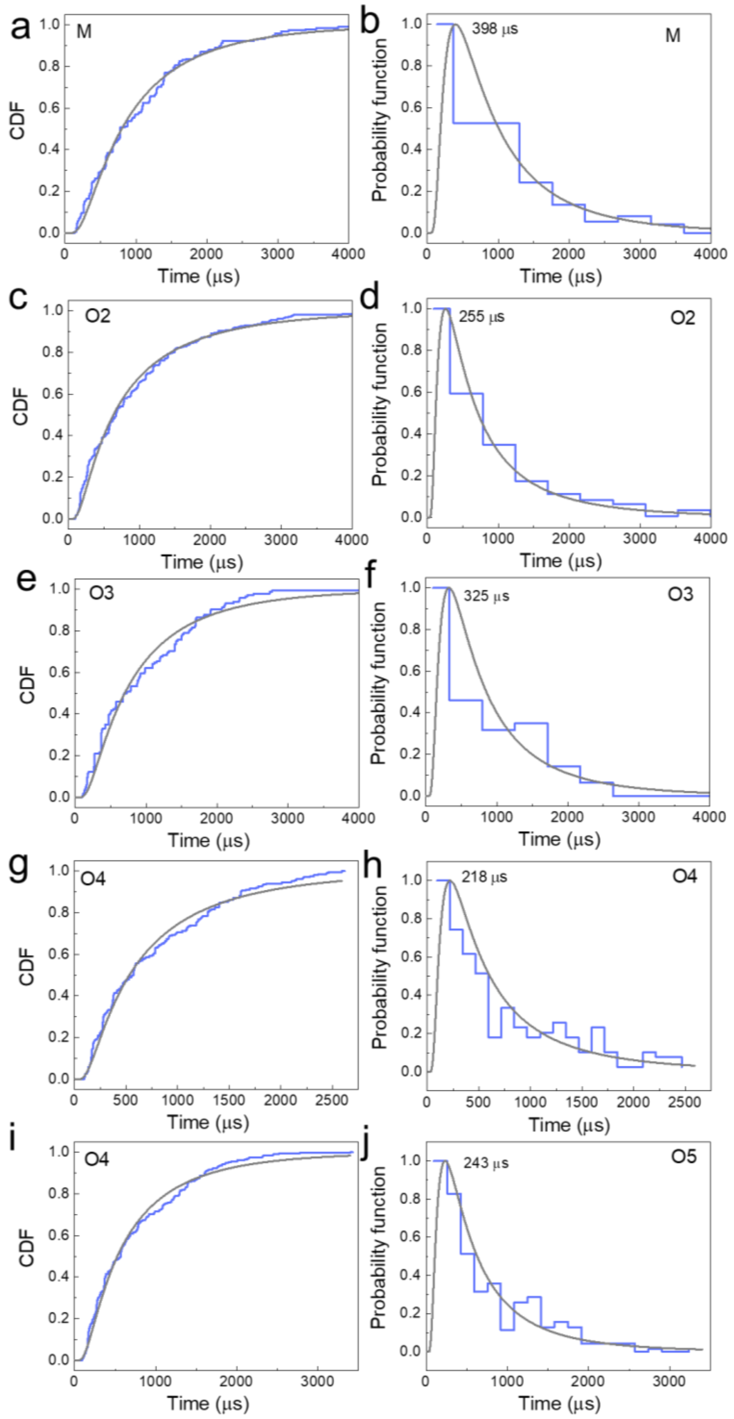


**Supplementary Figure S12. Cumulative density functions (CDFs) and histograms with PDFs of dwell times $t_d$ of experimentally measured resistive pulses from Tau monomers**

**and low-n oligomers**. (**a**, **b**) monomer (M), (**c**, **d**) dimer (O2), (**e**, **f**) trimer (O3), (**g**, **h**) tetramer (O4), (**i**, **j**) pentamer (O5). The distributions include the $t_d$ values of all detected resistive pulses that were longer than 20 μs and show that the most probable $t_d$ values for M, O1, O2, O3, O4, and O5 were 398 μs, 255 μs, 325 μs, 218 μs and 243 μs, respectively. The cumulative distribution functions (CDFs) were fitted to the experimental data and subsequently differentiated to obtain the corresponding probability density functions (PDFs, grey curves).

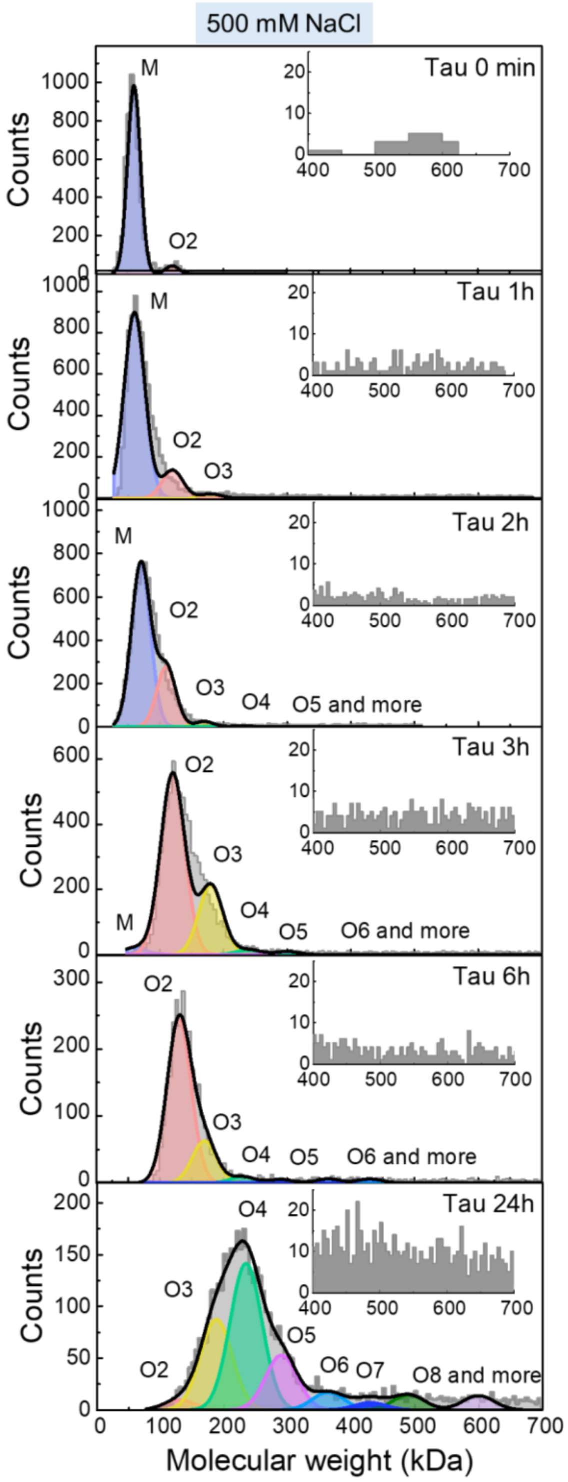

**Supplementary Figure S13. Mass photometry of Tau oligomers in the same high salt buffer as the one used for nanopore recordings.** Molecular weight (kDa) distributions of Tau oligomers with varying time points during protein aggregation in the same buffer as the one used for nanopore recordings with PLY pores containing 500 mM NaCl with 50 mM Tris-HCl, pH 7.5. The insets show a magnified y-axis of molecular weights ranging from 400 to 700 kDa, corresponding to higher-order Tau oligomers.

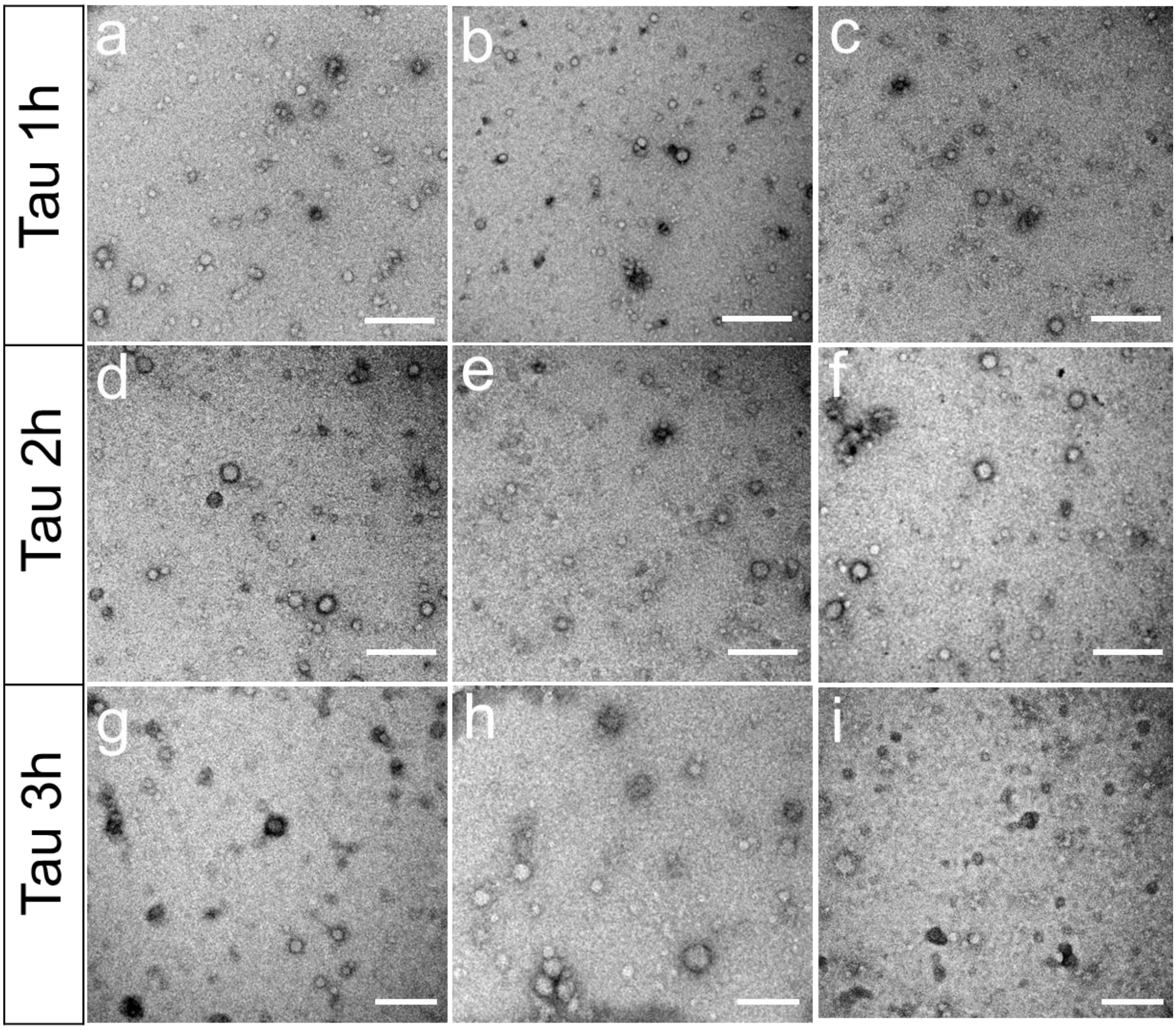


**Supplementary Figure S14. TEM micrographs of Tau oligomers after various durations of aggregation.** Representative images are shown for Tau samples incubated for (a–c) 1 h, (d–f) 2 h, and (g–i) 3 h. Scale bar: 200 nm.

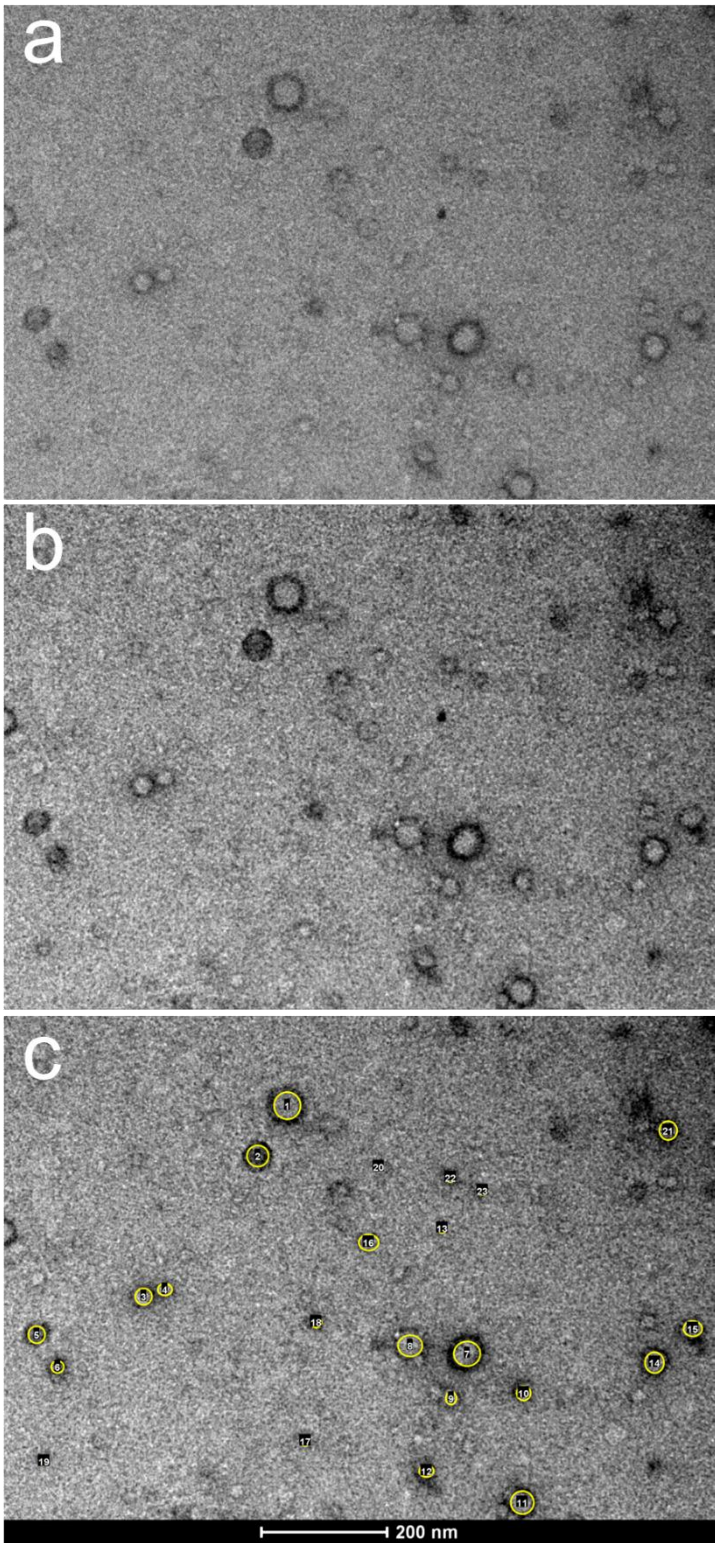


**Supplementary Figure S15. TEM image analysis of Tau oligomers.** (a–c) Representative steps of the image analysis workflow for Tau oligomers obtained from transmission electron microscopy (TEM) micrographs. (a) Raw TEM image of Tau oligomers. (b) Brightness and contrast–adjusted image after mean filtering (radius 2 pixels) to enhance particle visibility. (c) Image showing selected oligomers highlighted in yellow, used for size determination based on Feret's diameter measurements in ImageJ (Fiji).

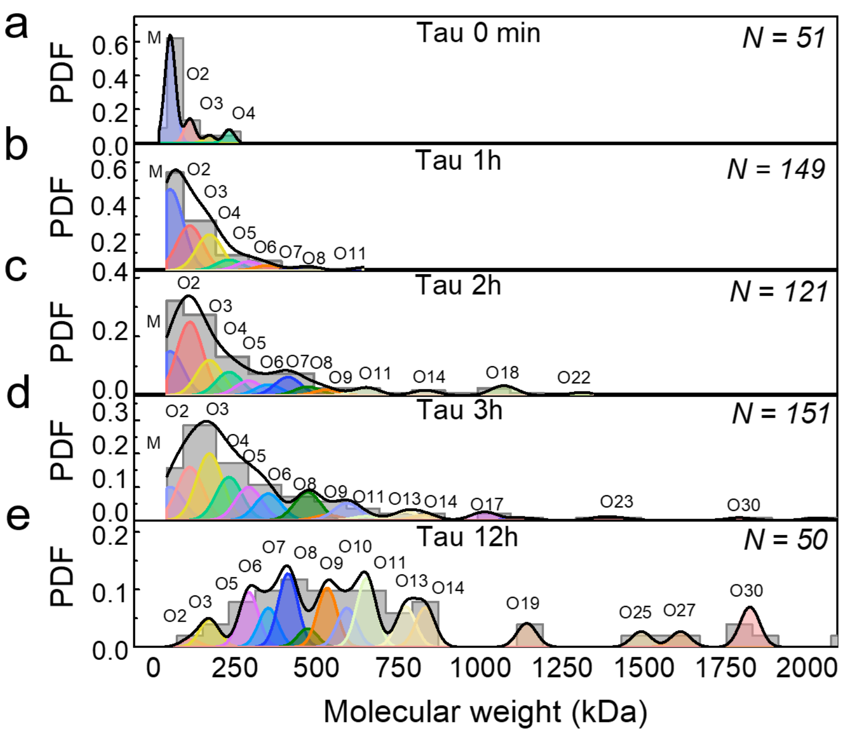


**Supplementary Figure S16. Molecular weight distribution of Tau oligomers (0 min, 1h, 2h, 3h and 12h) determined using TEM.** Different-sized oligomer subpopulations (as shown in **Figure 3e**) are marked as M to O30 based on molecular weight from monomer to 30-mer.

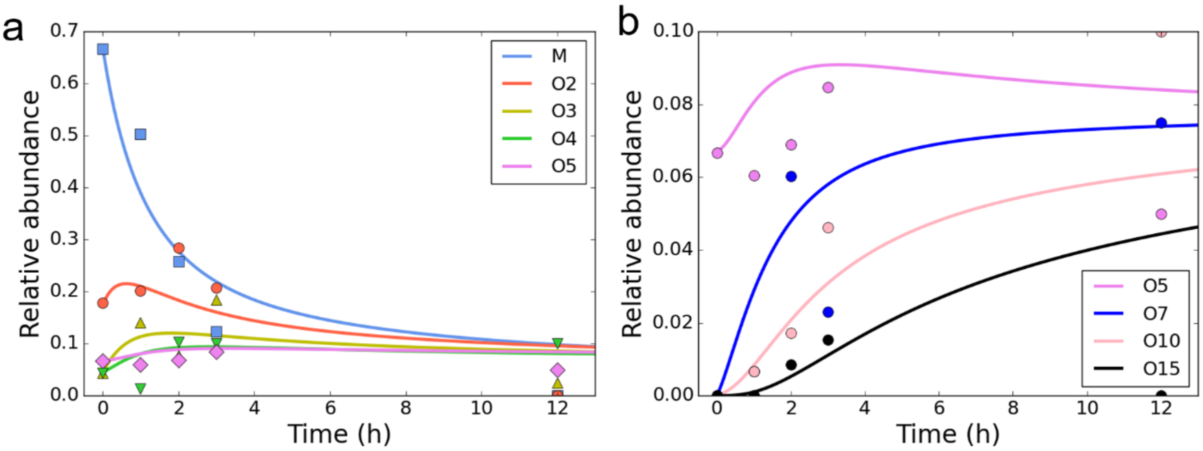


**Supplementary Figure S17. Time evolution of the abundance of various subpopulations of Tau oligomers determined using TEM.** Relative abundances of Tau monomers (M) and oligomers comprising 2–5 monomeric units (O2–O5) (a) and 5, 7, 10, and 15 monomeric units (O5, O7, O10, and O15) (b) as determined from TEM images acquired at different incubation times. The continuous lines are fits to the experimentally determined relative abundances using the colloidal aggregation model described in **Supplementary Note 7**. The resulting activation barrier was $10.7\ k_BT$, in close agreement with the value of $10.5\ k_BT$,obtained from the nanopore measurements.

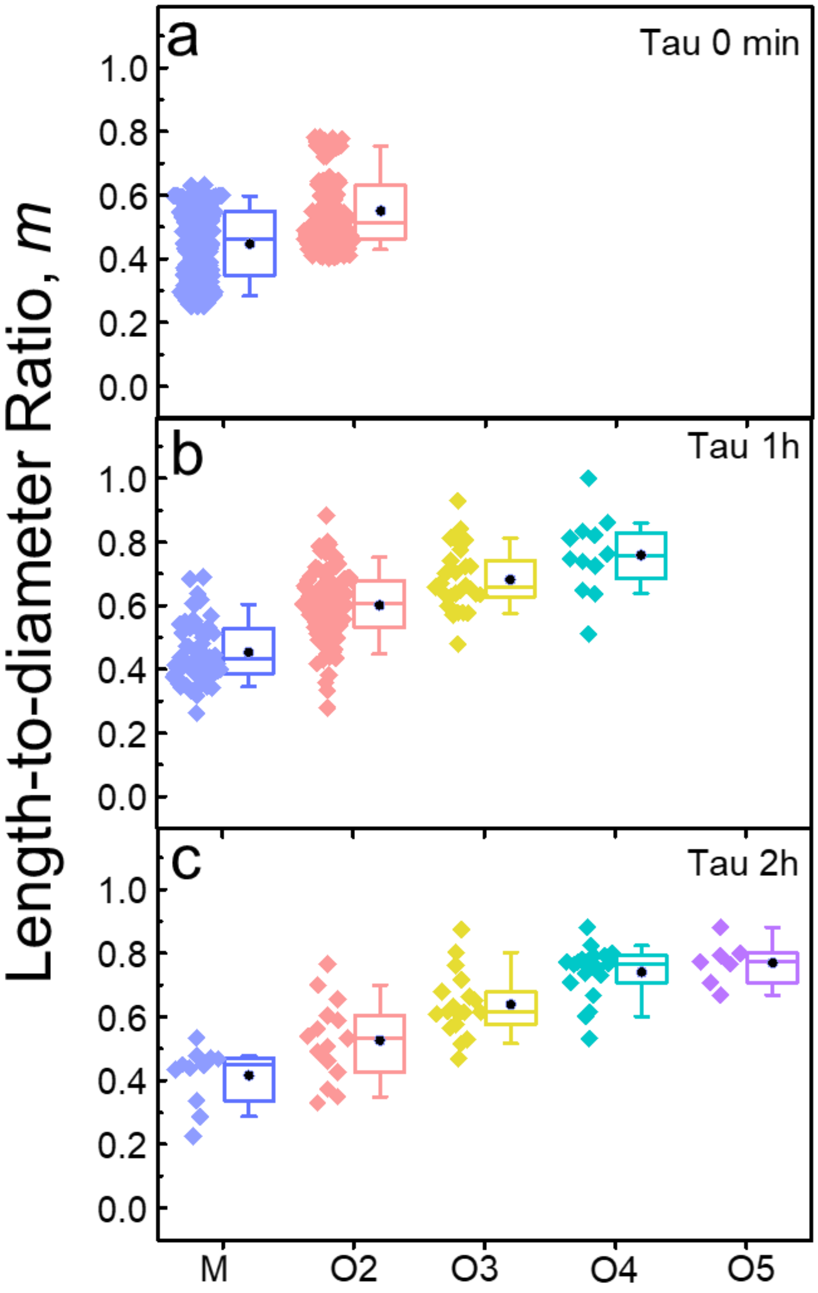


**Supplementary Figure S18. Single-molecule shape estimation of Tau oligomers in solution.** (**a-c**) Comparison of length-to-diameter ratios ($m$) determined from resistive pulses of Tau oligomer samples using PLY nanopores and a recording buffer containing 500 mM NaCl, Tris-HCl, pH 7.5. Measured length-to-diameter ratio ($m$; **Figure 4**) of Tau monomer and oligomers during aggregation after various time points (0 to 2h) as determined by PLY nanopores.

### Supplementary Table

**Supplementary Table S1. Estimation of reference values of excluded volumes and length-to-diameter ratios using molecular weight (MW)[8] of the protein or different models for approximating ellipsoidal shape.**[1,15]

| **Analyte protein** | **Reference excluded volume (Λ ; nm³) and length-to-diameter (*m*)** | | |
|---|---|---|---|
| | using MW of protein Ref [7] | data analysis package used in this study Ref [1] | data analysis package Ref [14] |
| Fab | Λ ($nm^3$) = 54 | Λ ($nm^3$) = 69<br>*m* = 0.6 | Λ ($nm^3$) = 81.2<br>*m* = 0.67 |
| ConA dimer | Λ ($nm^3$) = 64 | Λ ($nm^3$) = 98.8<br>*m* = 1.52 | Λ ($nm^3$) = 105.3<br>*m* = 1.66 |
| Hb | Λ ($nm^3$) = 78 | Λ ($nm^3$) = 115.1<br>*m* = 0.82 | Λ ($nm^3$) = 118.6<br>*m* = 0.85 |
| HSA | Λ ($nm^3$) = 80 | Λ ($nm^3$) = 105.3<br>*m* = 1.24 | Λ ($nm^3$) = 115.0<br>*m* =1.42 |
| ConA tetramer | Λ ($nm^3$) = 125 | Λ ($nm^3$) = 187.6<br>*m* = 0.87 | Λ ($nm^3$) = 182.1<br>*m* =0.88 |

### *Supporting Information references:*


1. Chanakul, W. *et al.* Large and Stable Nanopores Formed by Complement Component 9 for Characterizing Single Folded Proteins. *ACS Nano* **19**, 5240–5252 (2025).

2. Cruickshank, C. C., Minchin, R. F., Le Dain, A. C. & Martinac, B. Estimation of the pore size of the large-conductance mechanosensitive ion channel of Escherichia coli. *Biophys. J.* **73**, 1925–1931 (1997).

3. Fennouri, A. *et al.* Tuning the Diameter, Stability, and Membrane Affinity of Peptide Pores by DNA-Programmed Self-Assembly. *ACS Nano* **15**, 11263–11275 (2021).

4. van Pee, K. *et al.* CryoEM structures of membrane pore and prepore complex reveal cytolytic mechanism of Pneumolysin. *Elife* **6**, e23644 (2017).

5. van Pee, K., Mulvihill, E., Müller, D. J. & Yildiz, Ö. Unraveling the Pore-Forming Steps of Pneumolysin from *Streptococcus pneumoniae*. *Nano Lett.* **16**, 7915–7924 (2016).

6. Yusko, E. C. *et al.* Real-time shape approximation and fingerprinting of single proteins using a nanopore. *Nat. Nanotechnol.* **12**, 360–367 (2017).

7. Houghtaling, J. *et al.* Estimation of Shape, Volume, and Dipole Moment of Individual Proteins Freely Transiting a Synthetic Nanopore. *ACS Nano* **13**, 5231–5242 (2019).

8. Erickson, H. P. Size and Shape of Protein Molecules at the Nanometer Level Determined by Sedimentation, Gel Filtration, and Electron Microscopy. *Biol. Proced. Online* **11**, 32–51 (2009).

9. De Carlo, S. & Harris, J. R. Negative staining and cryo-negative staining of macromolecules and viruses for TEM. *Micron* **42**, 117–131 (2011).

10. Kayed, R. *et al.* Common Structure of Soluble Amyloid Oligomers Implies Common Mechanism of Pathogenesis. *Science (1979).* **300**, 486–489 (2003).

11. Lashuel, H. A. *et al.* α-Synuclein, Especially the Parkinson's Disease-associated Mutants, Forms Pore-like Annular and Tubular Protofibrils. *J. Mol. Biol.* **322**, 1089–1102 (2002).

12. Tian, Y., Liang, R., Kumar, A., Szwedziak, P. & Viles, J. H. 3D-visualization of amyloid-β oligomer interactions with lipid membranes by cryo-electron tomography. *Chem. Sci.* **12**, 6896–6907 (2021).

13. Viles, J. H. Imaging Amyloid-β Membrane Interactions: Ion-Channel Pores and Lipid-Bilayer Permeability in Alzheimer's Disease. *Angew. Chem. Int. Ed.* **62**, e202215785 (2023).

14. Atkins, P. W. . *Physical Chemistry : A Very Short Introduction*. (Oxford University Press, 2014).

15. Li, Y., Ying, C. & Mayer, M. Continuous, Low Latency Estimation of the Size and Shape of Single Proteins from Real-Time Nanopore Data. *Anal. Chem.* **98**, 224–234 (2026).
